# New evidence for the presence and function of phosphoinositides (PPIs) in the chloroplast

**DOI:** 10.1101/2024.09.06.611683

**Authors:** Mastoureh Sedaghatmehr, Frieda Rößler, Alexander P. Hertle

## Abstract

Essential proteins such as the chloroplast-localized Sec14-like protein (CPSFL1) and the vesicle-inducing protein in plastids (VIPP1) have a high affinity for binding phosphoinositides (PPIs) in vitro. PPIs are a specific class of phospholipids characterized by a phosphorylated inositol head group, and while they make up a small fraction of total phospholipids, they are crucial for various regulatory functions. However, the precise subcellular localization of most PPI species in plants remains unclear due to their rapid turnover and low abundance Currently there is no documented evidence for the presence and function of phosphoinositides (PPIs) in chloroplasts. In our study, we developed genetically encoded biosensors targeted to plastids, allowing for the detection of several PPI isoforms, including PI3P, PI4P, PI5P, PI(4,5)P_2_, and PI(3,5)P_2_ within chloroplasts. We demonstrated the specificity of these biosensors through immunological methods and observed changes in their distribution patterns when co-expressed with PPI-modifying enzymes (cTP-SAC7, cTP-PTEN, and cTP-dOCRL). Our findings also revealed the association and potential interaction between PI3P and VIPP1. Importantly, we found that elevated PPI levels during stress conditions led to altered biosensor localization, and plants expressing PPI modifiers showed increased sensitivity to drought stress, highlighting the role of PPIs in plant stress responses.

## Introduction

Phosphatidylinositols (PtdIns or PIs) are low-abundance lipids with significant regulatory functions, governing a wide range of physiological processes in eukaryotes. PtdIns are produced in the endoplasmic reticulum (ER) and subsequently transported to various subcellular sites through a combination of lipid transfer proteins and vesicle trafficking mechanisms (Phillips et al., 2006; Heilmann, 2009). Phosphorylated derivatives of PtdIns are generated by a group of enzymes, including phosphatidylinositol kinases (PIKs) and phosphatidylinositol phosphatases (Mueller-Roeber and Pical, 2002). PIK enzymes phosphorylate three out of the five available hydroxyl groups on the inositol ring, specifically at positions 3′, 4′, and 5′. This process generates seven phosphatidylinositol phosphates (PtdInsPs or PPIs) consisting of three monophosphates (PI3P, PI4P, and PI5P), three bisphosphates PI(3,4)P_2_, PI(4,5)P_2_, and PI(3,5)P_2_ and one triphosphate PI(3,4,5)P_3_. Among them, the presence of PI(3,4)P_2_ and PI(3,4,5)P_3_ has not yet been confirmed in plants (Mueller-Roeber and Pical, 2002; Zarreen et al., 2023). To date, no homologs of the heterodimeric inositol lipid 3-kinases have been identified in plant cells to phosphorylate PI4P to PI(3,4)P_2_ and PI(4,5)P_2_ to PI(3,4,5)P_3_. PI5P is found in plants, yet an enzyme responsible for its production remains unidentified (Mueller-Roeber and Pical, 2002; Heilmann, 2009; Zarreen et al., 2023).

PPIs are being recognized as new secondary messengers within plant cells. Besides their crucial role under favorable conditions, they are quickly generated when plants are exposed to a wide range of environmental stresses, indicating their importance under stress conditions (Munnik and Meijer, 2001; Meijer and Munnik, 2003; Wang et al., 2008; Xue et al., 2009; Munnik and Vermeer, 2010; Pribat et al., 2011; Zhai et al., 2013; Zarreen et al., 2023). PI3P, PI5P, PI(3,5)P_2_, and PI(4,5)P_2_ exhibited increased levels in response to salt and osmotic stress, likely due to the induction of relevant PI kinases or the action of lipid phosphatases, resulting from the degradation products of other PPIs. (Meijer et al., 1999; Pical et al., 1999; DeWald et al., 2001; Meijer et al., 2001; Meijer and Munnik, 2003, König et al., 2008). Individual studies have reported alteration in PPI kinase and phosphatase activity under stress conditions. Salt and osmotic stress have been shown to boost both PI 3-kinase and PI3P 5-kinase activities in plants. (Meijer et al., 1999; Meijer et al., 2001; Munnik and Meijer, 2001; Leshem et al., 2007, König et al., 2008). PTEN (phosphatase and tensin homolog deleted on chromosome ten) proteins in vascular plants play a crucial role in regulating 3-phosphorylated PI signaling events. The transcript levels of *AtPTEN2* exhibited changes in response to various stresses, including conditions of salt and osmotic stress (Pribat et al., 2011). Similarly, environmental stresses such as cold, salt, and drought stimulate the expression and activity of phosphoinositide-specific phospholipase C (PLC) (Hirayama et al., 1995; Munnik and Meijer, 2001). PLC induces the hydrolysis of PI(4,5)P_2_ to generate the two signaling molecules, inositol-_1,4,5_-trisphosphate (InsP_3_) and diacylglycerol (DAG) (Hirayama et al., 1995; Testerink and Munnik, 2011; Zarreen et al., 2023). PLC has the ability to enhance salt tolerance in rice as well as drought tolerance in maize and tobacco (Wang et al., 2008; Zhai et al., 2013; Li et al., 2017). The expression of the well-characterized PI4K kinase, *AtPI4Kγ3*, is induced upon exposure to salt stress. *Arabidopsis* plants overexpressing PI4Kγ3 exhibit increased stress tolerance and inducible expression of stress-responsive genes (Akhter et al., 2016). All these investigations suggest that alterations in PPI concentration are tightly regulated and play a critical role in facilitating precise PPI signaling.

The unique characteristics of PPIs, including their low abundance, rapid turnover, and diversity, make them dynamic signaling molecules. PPIs have the ability to bind to specific proteins, which can result in changes in the proteins’ localization, conformation, and activity. Additionally, PPIs can influence membrane curvature. These capabilities are crucial aspects of PPI function in facilitating downstream signaling in all eukaryotes (Meijer and Munnik, 2003; Xue et al., 2009; Munnik and Vermeer, 2010).

For instance, PPI affinity chromatography and mass spectrometry analysis confirmed that PI(3,5)P_2_ and PI3P bind to ROF1 (ROTAMASE FKBP1) in Arabidopsis (Karali et al., 2012). ROF1 is associated with responses to heat, salt, and osmotic stress (Meiri and Breiman, 2009; Karali et al., 2012). The authors suggested that the interaction of ROF1 with PPI causes a change in its localization under salt stress (Karali et al., 2012). Similarly, PI(4,5)P_2_ plays a role in regulating ion channels and proteins associated with cytoskeletal modification and membrane trafficking (McLaughlin et al., 2002; Suh et al., 2006; Munnik and Vermeer, 2010). While the functions of PtdIns in membrane trafficking, endomembrane organization, and cytoskeletal rearrangement are well-established, their pivotal roles in signaling and their subcellular localization are only emerging in plants. Due to their rapid turnover and limited abundance, the precise subcellular localization of most PPI species in plants remains incomplete. The existence and function of PPI in the nucleus of animals, yeast, and plant cells has been documented (Horton et al., 2007; Dieck et al., 2012; Gerth et al., 2017). It is widely recognized that nuclear PtdIns play a significant role in regulating various processes, including DNA replication, chromatin remodeling, transcription, stress responses, and hormone signaling in plant, animal, and yeast cells (Dieck et al., 2012; Zarreen et al., 2023). The unique role of PPIs in the nucleus, distinct from their function in the plasma membrane, highlights the existence of distinct PPI signaling pathways within various cellular organelles.

A recent biochemical study has highlighted the critical role of PI4P in the functioning of another cellular organelle, chloroplasts. They revealed that inhibiting PI4Ks through chemical inhibitor treatment in *Arabidopsis* seedlings accelerates chloroplast division. It has also been reported that PI4P specifically binds to PLASTID DIVISION1 (PDV1) and PDV2, exerting a negative regulatory effect on chloroplast division. Of note, PDV1 and PDV2 proteins are located at the outer envelope membrane of the chloroplast, and the PI4P lipid-signaling pathway, involved in chloroplast division via PDVs takes place at the outer envelope membrane. Therefore, the presence and function of PPIs inside the chloroplasts remain unexplored thus far (Okazaki et al., 2015).

Lipid transport across the envelope membranes is highly controlled and restricted (Karki et al., 2019). Furthermore, the enzymes responsible for PPI biosynthesis in plastids have so far not been identified. This makes the presence of these “eukaryotic” lipids in the inner chloroplast membranes (e.g. inner envelope and thylakoids) rather unlikely. Nevertheless, experiments using radioactive labeling with ATP, as well as mass spectrometric analysis of plastidial endomembranes, have detected low amounts of individual PPI species (König et al., 2008; Siegenthaler et al., 1997). Contamination through membrane contacts with the ER or thylakoid/envelope membranes would be one explanation. Yet, the presence and function of PPIs in plastids cannot be denied either. Additionally, studies on membrane remodeling protein VIPP1 and chloroplast vesicle traffic regulator CPSFL1 support the latter by to PPIs with high specificity *in vitro* (Theis et al.2019, Hertle et al., 2020)

In this study, our results present evidence for the presence of PPIs within the chloroplasts of tobacco, as indicated by various PPI biosensors and immunodetection. We observed rapid relocation of PPI biosensors within the chloroplasts of *Arabidopsis* under abiotic stress conditions. Additionally, plants expressing chloroplast-targeted PPI modifiers exhibited increased sensitivity to drought and heat stress. Overall, our investigation sheds light on the existence of PPIs in the chloroplast and their potential involvement in the PPI signaling pathway during stress conditions.

## Results

### Generation and sub-organelle localization of a set of PPI biosensors in chloroplasts

Monitoring PPI poses a consistent challenge due to the rapid turnover, tight regulation, and low quantities of PPIs present within all living cells (Meijer and Munnik, 2003; Jensen et al., 2022). Indirect evidence indicating the presence of PPIs in the chloroplast is mainly limited to *in vitro* studies (Gou et al., 2015; Okazaki et al., 2015; Theis et al., 2019; Hertle et al., 2020). Nevertheless, a current research gap exists concerning the *in vivo* study of PPIs within chloroplasts.

Genetically encoded biosensors have been widely employed to study the distribution of PPIs in functional living cells (Varnai and Balla, 2006; Balla, 2013; Simon et al., 2014). Biosensors consist of lipid-binding domains that interact specifically with known PPIs and can be easily tracked by fusion with a fluorescent protein. To investigate the presence of various PPIs in the lipid composition of chloroplasts, we generated a collection of biosensors that contain a chloroplast transit peptide (cTP), yellow fluorescence protein or mCherry, and a lipid-binding domain. To individually and specifically detect PI3P, PI4P, PI5P, PI(4,5)P_2_, and PI(3,5)P_2_, the coding sequences of the following domains were used. The pleckstrin homology (PH) domain of human ARHGEF9 was used for PI5P binding. The PH domain of human Rho guanine nucleotide exchange factor 7 (ARHGEF7) was used for PI(3,5)P_2_ detection (Singh et al., 2021). The PH domain of rat four-phosphate adaptor protein 1 (FAPP1) was used for PI4P detection. The PH domain of mouse 1-phosphatidylinositol-4,5-bisphosphate diesterase delta (Plcd1) was used for PI(4,5)P_2_ detection. The FYVE domain of human hepatocyte growth factor-regulated tyrosine kinase substrate (HGS11) was used for PI3P detection (Simon et al., 2014). These domains were codon-optimized for expression in *Arabidopsis* and fused to the C-terminal region of a chloroplast-targeted fluorescent reporter (Fig. 1a). Whereas some PH domains bind to more than one PPIs isoform, the employed domains for PI3P, PI4P and PI(4,5)P_2_ detection have been shown in numerous *in vivo* studies in plants to be specific for their substrate (Varnai and Balla, 2006; Balla, 2013; Simon et al., 2014). Moreover, specific recognition of PI5P and PI(3,5)P_2_ by the lipid binding domains of ARHGEF7 and 9 has been shown *in vitro* (Singh et al.2021).

**Figure 1:**
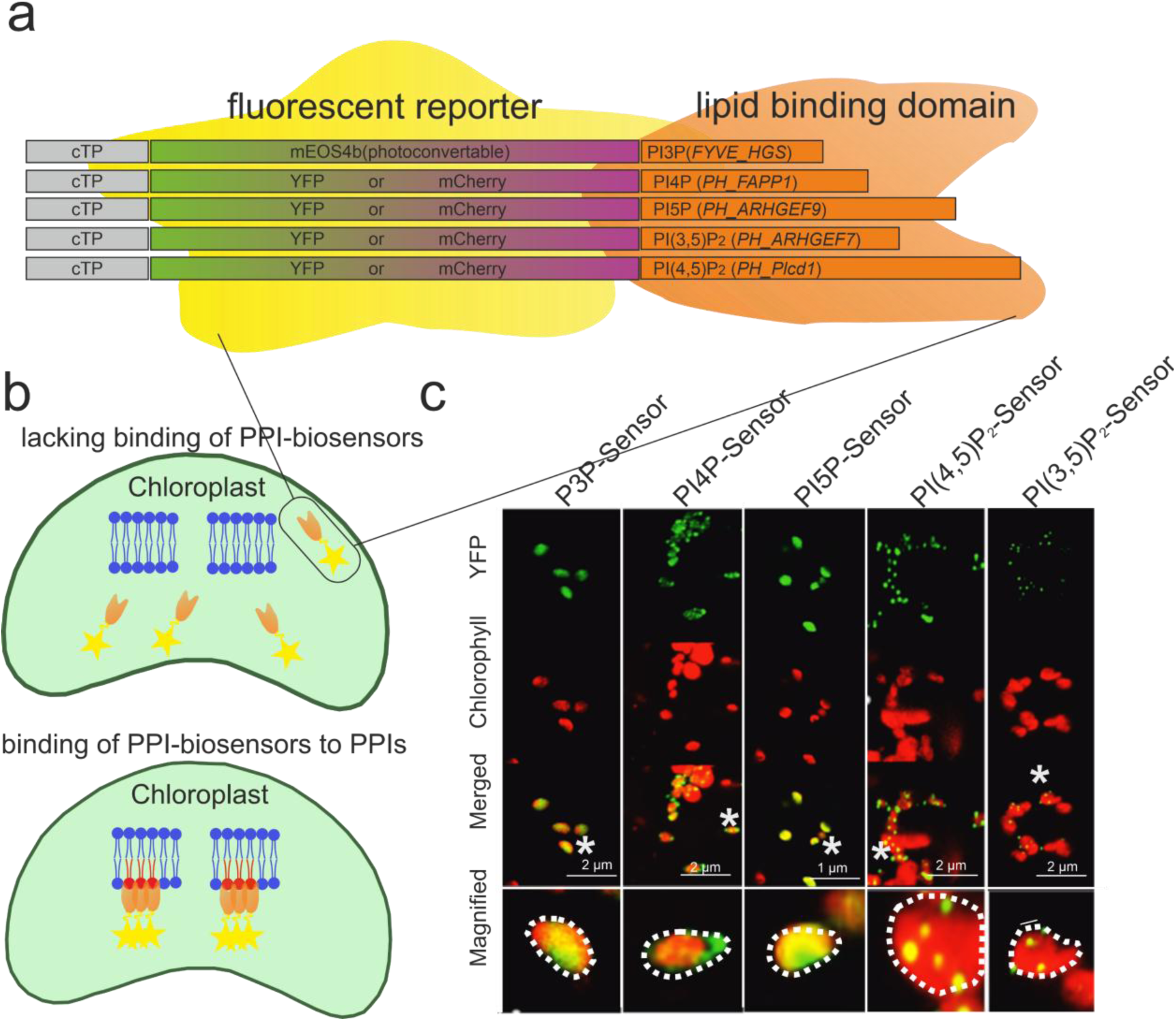
Expression of YFP-PI3P, YFP-PI4P, YFP-PI5P, YFP-PI(4,5)P_2_ and YFP-PI(3,5)P_2_ biosensors in chloroplasts of tobacco and identification of their sub-organelle distribution. A. The collection of biosensors is constructed to detect the different PPIs individually. Each specific biosensor consists of a chloroplast transit peptide and a fusion between lipid-binding domains that interacts specifically with each PPI and a yellow fluorescent protein (YFP), mCherry or mEOS4b. B. scheme representing the distribution pattern of PPI biosensors in chloroplast with or without binding to specific PPIs. The upper panel represents a lack of binding of the PPI biosensor with diffuse localization of PPI biosensors in the chloroplast. Lower panel represents the binding of PPI biosensors to PPI-containing membranes with distinct localization of PPI biosensors. C. Confocal images of tobacco leaves transiently expressing PPI biosensors in the chloroplasts. Lacking the binding of the PI3P and PI5P biosensors results in diffuse localization of mEOS4b-PI3P and YFP-PI5P biosensors in the chloroplast. The binding of YFP-PI4P, YFP-PI(4,5)P_2_ and YFP-PI(3,5)P_2_ biosensors results in distinct punctate localization of PI biosensors indicating substrate recognition.

To visualize their distribution pattern in plastids, we performed a transient expression of each of the designed biosensor constructs in tobacco leaves using infiltration with transgenic *Agrobacteria* to initiate transient expression (shown in Fig. **1a**). YFP fluorescence was observed 48 hours after Agrobacterium infiltration in the chloroplast of all transformed cells due to the presence of chloroplast transit peptide. We further analyzed the sub-organelle distribution of PPI biosensors in the chloroplasts. We observed a diffuse localization of YFP-PI3P and YFP-PI5P biosensors within the chloroplast. In contrast, binding of YFP-PI4P, YFP-PI(4,5)P_2_, and YFP-PI(3,5)P_2_ biosensors to PPI-containing membrane leads to a different localization reflected in distinct punctate structures within or surrounding the thylakoids represented by chlorophyll autofluorescence of chloroplasts (Fig. **1b, c**). Counting puncta in several chloroplasts gave a constant and characteristic number of puncta for respective sensors. Furthermore, PI(3,5)P_2_ and PI(4,5)P_2_ biosensors, with a lower number of puncta colocalized with the chlorophyll fluorescence, whereas PI4P sensors with a much higher number of smaller puncta localized differently in a peripheral region of the chloroplast surrounding the thylakoids as indicated by chlorophyll fluorescence (Supporting Information Fig. S1 a, b).

The presence of PI4P in the outer envelope of the chloroplast has been previously demonstrated (Okazaki et al., 2015). To confirm that the YFP-PI4P biosensors detect their substrate within the chloroplast and not at the outer envelope of the chloroplast, we treated isolated chloroplasts from plants expressing YFP-PI4P biosensor with the protease thermolysin. Thermolysin effectively breaks down proteins located on the surface of chloroplasts, but it does not penetrate the intermembrane space (Zufferey et al., 2017). Chloroplasts subjected to thermolysin treatment were subsequently analyzed by SDS-PAGE and subjected to immunoblot analysis. TOC159, a resident of the outer chloroplast envelope membrane and known to be thermolysin-sensitive was notably reduced following thermolysin treatment. In contrast, the YFP-PI4P biosensor protein remained largely unaffected by the treatment (Supporting Information Fig. S1c). This observation indicates that the YFP-PI4P biosensor is mainly associated with membranes within the chloroplasts and is not readily accessible for thermolysin-mediated degradation at the outer surface membrane. To further support the presence of PI4P within the inner membranes of chloroplasts as well as its specificity, we performed fat blot assays using lipid standards together with membrane extracts and detected PI4P immunologically using PI4P specific antibodies (Weber et al., 2006) (Supporting Information Fig 1d). The same antibodies also recognized lipid extracts from chloroplasts and thylakoids isolated from *Arabidopsis* (Supporting Information Fig S1d).

### Chloroplast localized-PPI biosensors change their distribution upon PPI phosphatase expression

To further support the specificity of our PPI biosensors, we verified the sub-organelle distribution pattern of PPI biosensors in the chloroplasts of tobacco plants in the presence of PPI modifiers. For this purpose, we generated three distinct PPI phosphatases that feature a chloroplast transit peptide. These phosphatases include SUPPRESSOR OF ACTIN 7 (SAC7) phosphatase and tensin homolog deleted on chromosome ten (PTEN), and the phosphatase domain of *Drosophila melanogaster* Oculocerebrorenal Syndrome of Lowe protein (dOCRL) (Goulden et al., 2019; Thole et al., 2008; Doumane et al., 2021). SAC7, PTEN, and dOCRL can reversibly dephosphorylate PI4P, PI3P, and PI5P respectively, by their action as lipid phosphatases (Fig. **2a**). Co-expression of the PI(4,5)P_2_ biosensor with *cTP-SAC7* and *cTP-dOCRL*, modifiers specific to PI4P and PI5P, respectively, resulted in a significant and substantial alteration in the distribution pattern of the PI(4,5)P_2_ biosensor. The distinctive punctate structures of the PI(4,5)P_2_ biosensor were notably reduced and transformed into diffuse fluorophore signals in the presence of the SAC7 and dOCRL phosphatases (Fig. **2b, c**). In contrast, the punctate-like structures of the PI(4,5)P_2_ biosensor remained unchanged when co-expressed with the PI3P modifier, *cTP-PTEN* (Fig. **2c**). The punctate signals of the PI(4,5)P_2_ biosensor within the chloroplasts was also predominantly converted into diffuse fluorophore signals in the presence of the PI4P modifier SAC7. Since PI3P biosensors were equally distributed when targeted to plastids, co-expression of PTEN or SAC7 did not show any alternation (Fig. **2d**). Observed changes were quantified and analyzed for statistical relevance (Supporting Information Fig. S2a). A significant reduction of fluorescent puncta was observed for PI4,5P_2_ sensors upon dOCRL expression. Moreover, co-expression of SAC7 and PI4P sensors resulted in a significant reduction of the fluorescent puncta but an increase in a diffuse fluorescent signal.

**Figure 2:**
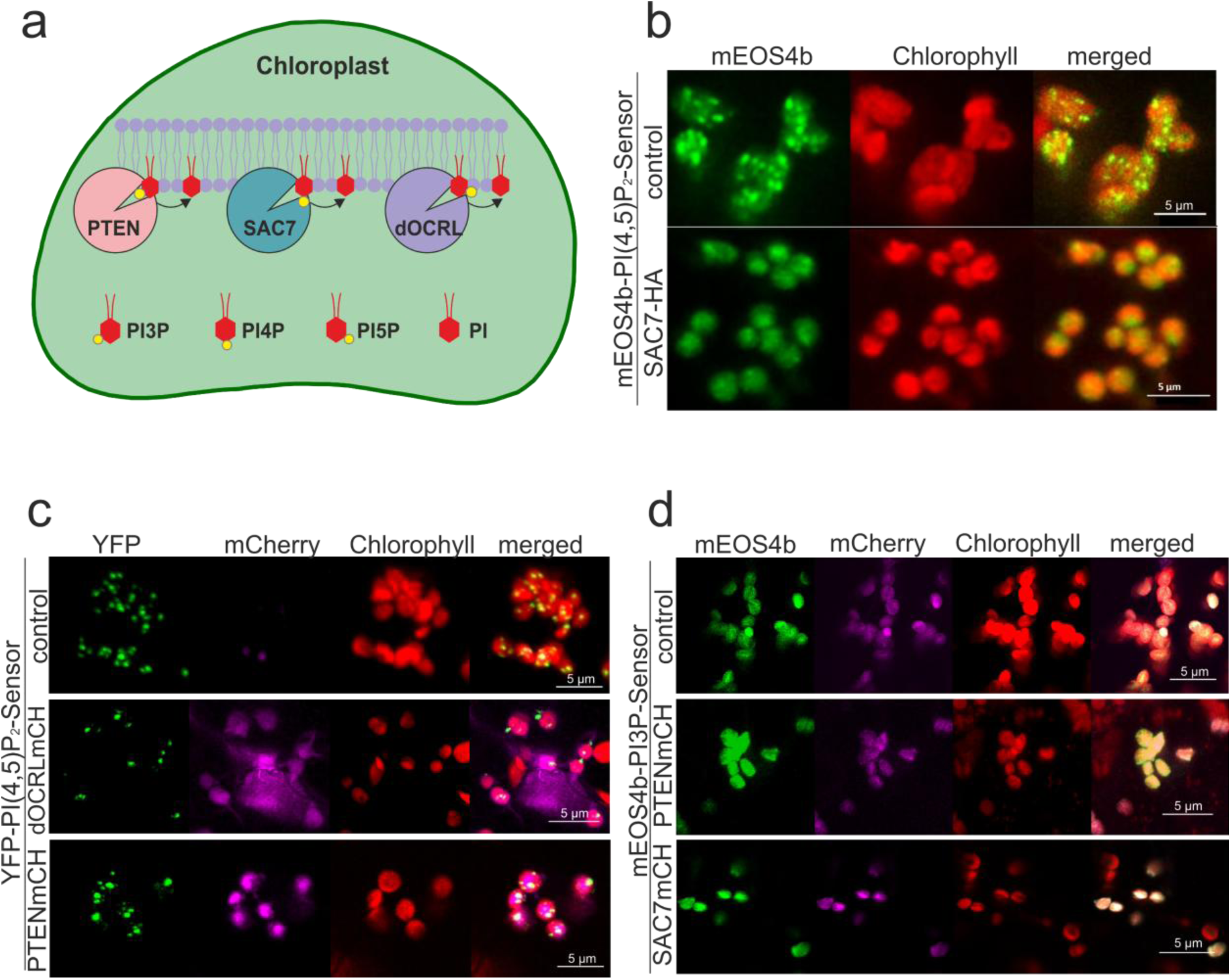
Sub-organelle distribution pattern of mEOS4b-PI3P, mEOS4b-PI4P and YFP-PI(4,5)P_2_ biosensors in the presence of the different PPI phosphatases in the chloroplasts. A. The illustration shows a simplified view of the phosphatase activity of PTEN, SAC7, and dOCRL in chloroplasts. B. Distribution pattern of mEOS4b-PI4P biosensors in the presence of phosphatase, SAC7, in the chloroplasts of transiently expressing tobacco. C. Distribution pattern YFP-PI(4,5)P_2_ biosensors without and in the presence of presence of PI5P and PI3P phosphatases, dOCRL-mCherry and PTEN-mCherry in the chloroplasts of tobacco. D. Distribution pattern of PI3P biosensors in the presence of PI3P and PI4P phosphatases, PTEN-mCherry and dOCRL-mCherry in the chloroplasts of tobacco. Fluorescence signals were visualized in tobacco leaves co-expressing PPI biosensors and phosphatases (as indicated) by fluorescence confocal microscopy.

We also examined the distribution of the same PI(4,5)P_2_ biosensor tagged with different fluorophores following co-expression. As expected, biosensors with identical substrate, YFP-PI(4,5)P_2_ and mCherry-PI(4,5)P_2_, co-localized with the same fluorescence pattern in the chloroplasts (Supporting Information Fig. S2b). Next, we co-expressed biosensors sensitive for PI(3,5)P_2_ tagged with YFP and a biosensor specific for PI4P and tagged with mCherry transiently in tobacco leaves. The sensors differed in their sub-organelle localization and in respect to the chlorophyll autofluorescence. In summary, these observations collectively demonstrate the specificity of our PPI biosensors in selectively detecting individual PPIs within the chloroplasts (Supporting Information Fig. S2c, d).

### VIPP1 may colocalize with PI3P rich membranes

To date, our understanding of the interaction between chloroplast proteins and PPIs has primarily relied on *in vitro* studies employing protein-lipid blot overlay assays (Gou et al., 2015; Okazaki et al., 2015; Theis et al., 2019; Hertle et al., 2020). We further assayed the colocalization of the chloroplast protein, VESICLE-INDUCING PROTEIN IN PLASTIDS 1 (VIPP1) with individual PPI biosensors. *In vitro* studies have already demonstrated the robust binding of *Chlamydomonas reinhardtii* VIPP1 to liposomal membranes containing PI4P, PI(4,5)P_2_, and PI(3,4,5)P_3_ (Theis et al., 2019).

To assess the *in vivo* association of VIPP1 with PPIs, we conducted colocalization between VIPP1-CFP and PPI biosensors within the chloroplasts. Interestingly, the simultaneous expression of *VIPP1-CFP* and *cTP-mEOS4b-PI3P* led to a significant alteration in the distribution pattern of mEOS4b-PI3P, resulting in the formation of distinct dot-like structures (Figure 3A). Significantly, these dot-like structures of mEOS4b-PI3P exhibited co-localization with VIPP1-CFP (Fig. **3a**). Interestingly, the overrepresentation of VIPP1 did not alter the sub-organelle distribution pattern of the PI(4,5)P_2_ biosensor (Fig. **3a**).

**Figure 3:**
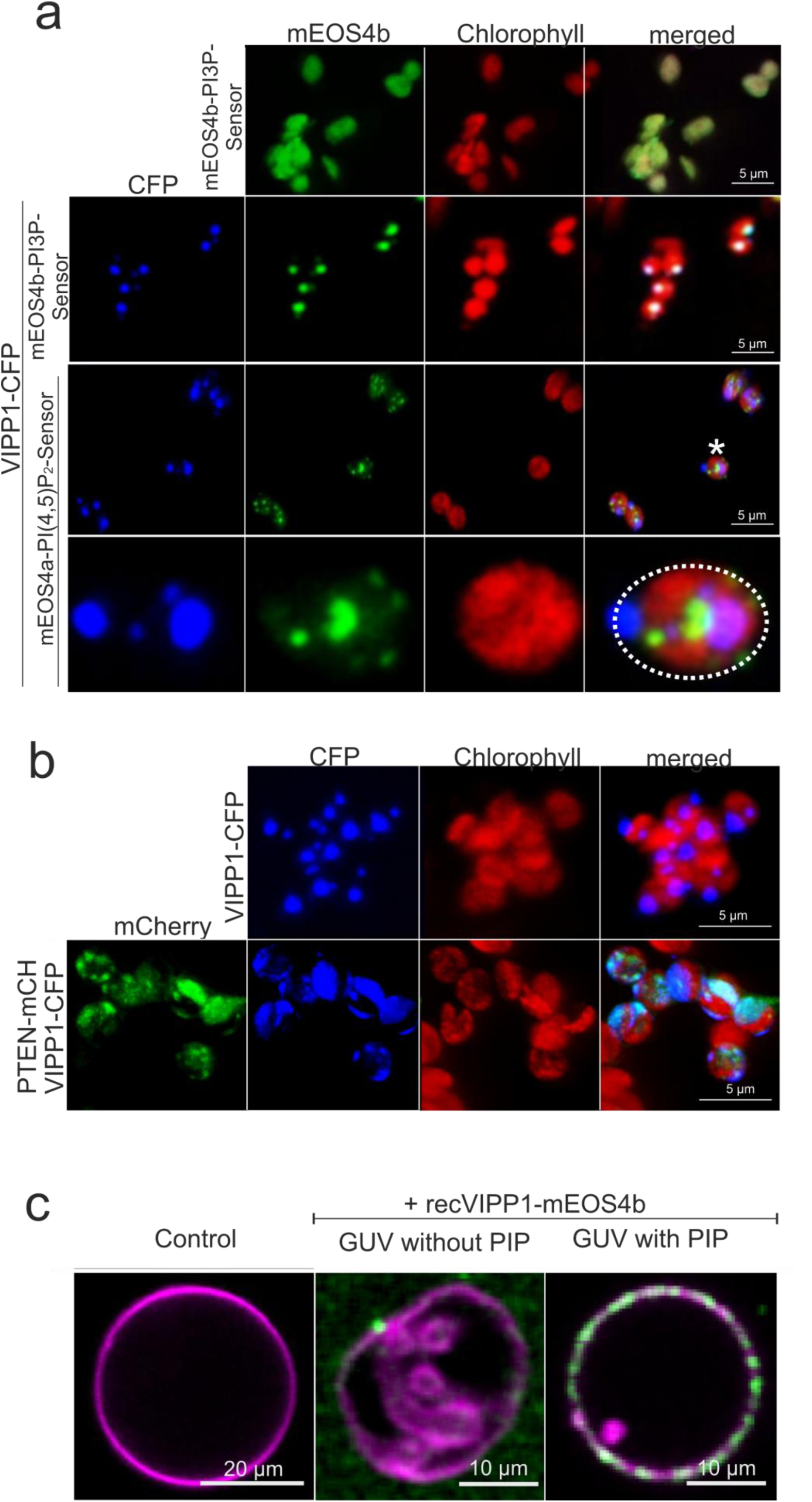
Co-localization of VIPP1 with cTP-mEOS4b-PI3P (FYVE) biosensor and its differential sub-organelle localization in the presence of the PtdIns3P phosphatase, PTEN in the chloroplasts. A. Confocal images of tobacco leaves transiently expressing mEOS4b-PI3P (FYVE) biosensor in the presence and absence of VIPP1-CFP (middle and upper panel). Middle panels show colocalization of mEOS4b-PI3P biosensor with VIPP1-CFP in the chloroplasts. Lower panel shows different localization of mEOS4b-PI(4,5)P_2_ biosensor and VIPP1-CFP in the chloroplasts. B. Sub-organelle distribution pattern of VIPP1-CFP in the presence of PI3P phosphatases, PTEN in the chloroplasts. C. Plant lipid based GUVs (magenta) with and without PI3P in the presence or absence of recombinant mEOS4b-VIPP1 (green).

Similarly, co-expression of the *VIPP1-CFP* with *cTP-mCherry-PTEN*, modifier specific to PI3P, resulted in a substantial alteration in the typical distribution pattern of VIPP1 within chloroplasts (Fig. **3b**). Subsequently, we examined whether the stability of VIPP1 and the configuration of its homooligomeric complexes were affected in the presence of PTEN. The amounts of VIPP1 remained unaltered in plants expressing *cTP-PTEN* (Supplementing Information Fig. S3a). To verify the complex state of VIPP1 in *cTP-PTEN*-expressing plants, thylakoid proteins from both WT and *cTP-PTEN* plants were subjected to treatment with Bis (Sulfosuccinimidyl) suberate (BS^3^). BS^3^ serves as a membrane-impermeable crosslinking agent to stabilize the complex state. Nevertheless, no significant differences were observed in terms of the multiple complex states of VIPP1 in *cTP-PTEN*-expressing plants (Supporting Information Fig. S3b). Our observation suggests that overexpression of VIPP1 may concentrate PI3P lipids in a specific location within the chloroplast, making them detectable by the PI3P biosensor, while in the absence of VIPP1 overexpression, the condensation of PI3P appears to be below the detection threshold of the PI3P biosensor (Fig. **1c**). Thus, we wanted to proof the sensitivity and specificity of the PI3P sensor by an independent approach. For this we purified chloroplasts and thylakoids purified form *Arabidopsis* and spinach respectively. Following lipid extraction, we performed fat blot assays together with a dilution series of PI3P lipid standards and probed them with PI3P specific FYVE-mEOS4b sensor proteins. Recombinant FYVE sensors recognized PI3P lipid standards as well as chloroplast and thylakoid lipid extracts from both species. Still, with a low concentration (<6.3 pmol) and obscured by a bright spot in the center of the lipid dot resembling the distribution of coextracted chlorophyll (Supporting Information Fig. S3c). Thus, we fractionated spinach chloroplasts and probed chlorophyll free inner envelope membranes, thylakoids and an array of chloroplast lipids to support specific recognition of PI3P sensor proteins. PI3P and inner chloroplast envelope membranes were recognized with high specificity (Supporting Information Fig. S3d).

The specificity of FYVE-mEOS4b sensors in the presence of thylakoid lipids and PI3P was further investigated by using recombinant sensor proteins and using giant unilammelar vesicles (GUVs) composed of MGDG, DGDG, SQDG, PG and PI in a stöchiometry of 52:26:6.5:9.5:1 (v/v) (Hölzl & Dörmann, 2019; LaBrant et al., 2018). Only PI3P containing GUVs were recognized by PI3P biosensor proteins (Supporting Information Fig. S3e). Next, we tested the binding of VIPP1 to plant lipid-based giant unilamellar vesicles (GUVs). While low binding of VIPP1 was observed in GUVs composed of MGDG, DGDG, PG, and SQDG, the inclusion of 0.1 mol% PI3P into the GUV lipid mixtures led to a pronounced recruitment of recombinant VIPP1 to the GUV surfaces in punctate pattern (Fig. **3c**). This punctate pattern of VIPP1could not be observed for FYVE sensor proteins which suggests a non-homogeneous distribution or local concentration of phosphatidylinositols (PPIs) induced by VIPP1 (Fig. **3a, c**).

Taken together, these results suggest the association and potential function of VIPP1 with PI3P *in vivo*.

### The localization of PPI sensors changes upon heat in chloroplasts of *Arabidopsis*

To investigate the presence and function of various PPIs in the chloroplasts, we generated transgenic *Arabidopsis* lines stably expressing our chloroplast localized biosensors (*cTP-YFP-PI4P*, *cTP-YFP-PI5P*, *cTP-YFP-PI(4,5)P_2_*, and *cTP-YFP-PI(3,5)P_2_*. Of note, none of the transgenic lines showed any visible developmental phenotypes. Next, we assessed the sub-chloroplasts distribution patterns of YFP-PI4P, YFP-PI5P, YFP-PI(4,5)P_2_, YFP-PI(3,5)P_2_ biosensors, and cTP-GFP within the chloroplasts of 10-day-old *Arabidopsis* seedlings under standard control conditions using confocal microscopy. We noted that all the PPI sensors displayed a diffuse localization within the chloroplast (Fig. **4a**). This observation suggests that the biosensors may not have encountered their intended targets for binding, or alternatively, the concentration of PPIs in the chloroplast of *Arabidopsis* seedlings might be below the detection limits of our designed PPI biosensors.

**Figure 4:**
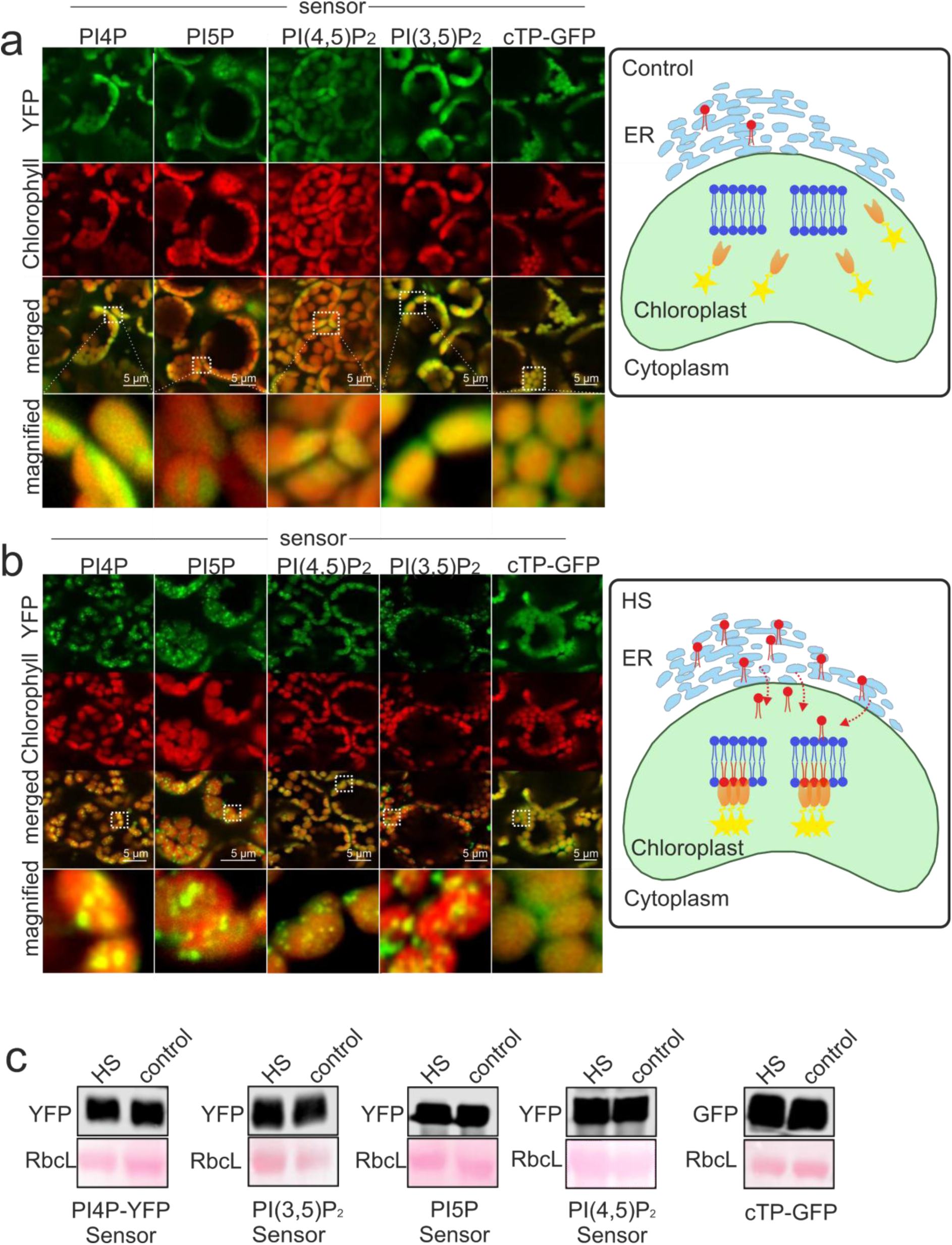
Sub-organelle distribution pattern of YFP-PI4P, YFP-PI5P, YFP-PI(4,5)P_2_ and YFP-PI(3,5)P_2_ biosensors as well as cTP-GFP in the chloroplast of *Arabidopsis* seedlings under control and heat stress (HS) conditions. A and B. Confocal images of 10-days-old *Arabidopsis* seedlings stably expressing *cTP-YFP-PI4P*, *cTP-YFP-PI5P*, *cTP-YFP-PI(4,5)P_2_* and c*TP-YFP-PI(3,5)P_2_* biosensors. (A) control and (B) HS (at 40°C for 30 minutes). The diagrams at the right represent PIP flux upon control and HS. C. Immunoblot analysis of cTP-GFP, YFP-PI4P, YFP-PI5P, YFP-PI(4,5)P and YFP-PI(3,5)P_2_ proteins under control and HS conditions using GFP antibody.

Furthermore, we decided to elevate the PPI flux within the cell, potentially including the chloroplast, and subsequently assess the sub-chloroplast distribution patterns of the PPI biosensors. Studies have demonstrated that the metabolism of PI and PPIs is significantly activated in response to heat stress in various organisms (Morales-Johansson et al., 2004; Lu et al., 2020) and diverse plant species (Mishkind et al., 2009; Annum et al., 2022; Zhu et al., 2022; Annum et al., 2023). To elevate the PPI flux, we subjected 10-day-old Arabidopsis seedlings to a 40°C temperature treatment for 30 minutes, after which we conducted confocal microscopy analysis. Surprisingly, we observed that all PPI biosensors re-localized within the chloroplasts in response to short heat stress treatment. As shown in Fig. **4b**, there was no change in the sub-chloroplast localization of GFP in plants expressing GFP targeted into chloroplasts. In contrast, we observed a clear translocation of sensors to PPI-containing membranes resulting in punctate-like structures in the plant expressing PPI biosensors (Fig. **4b**). Quantification of fluorescent loci in plastids containing PI4P, PI5P and PI(3,5)P_2_ sensor proteins showed similar numbers per chloroplast. Consistent with transient expression PI(4,5)P_2_ sensor containing plastids showed significantly fewer puncta as compared to other sensor lines (Supporting Information Fig. S4a, b). Although, expression of PPI biosensors is under the control of the 35S promoter, we additionally confirmed, through western blot analysis, that the heat stress treatment did not result in any noticeable changes in their abundance when compared to the control condition (Fig. **4c**). Moreover, we noticed that the distribution patterns of PI(4,5)P_2_-YFP biosensors remained unchanged in plastids of heat-stressed root cells, in comparison to untreated root cells (Supporting Information Fig. S4c). These results indicates that the relocation PI(4,5)P_2_ sensors observed in leaves is dependent on chloroplast development or thylakoid differentiation.

### Potential function of PPIs as a lipid signaling molecule in chloroplasts under abiotic stresses

Multiple studies have demonstrated stress-induced alterations in PI and PPI signaling, implying the pivotal roles of PPI signaling in plant adaptation to stress conditions (Hirayama et al., 1995; Meijer et al., 1999; Pical et al., 1999; DeWald et al., 2001; Meijer et al., 2001; Heilmann, 2009; Munnik and Vermeer, 2010; Zhu et al., 2022; Annum et al., 2023). We observed that almost all PPI biosensors re-localized within the chloroplasts in response to drought stress treatment, (Fig. **5a** and Supporting Information Fig. S5a) most probably due to increased PPI flux within the chloroplasts following drought stress. Quantification of fluorescent puncta per chloroplast showed a significant difference between PI4P and PI(4,5)P_2_ and PI(3,5)P_2_ lines. In contrast, no punctate structures were observed for chloroplasts of PI3P and PI5P lines (Supporting Information Fig. S5a). Next, to further explore the possible involvement of various PPIs within the chloroplasts in response to abiotic stress, we generated transgenic Arabidopsis lines that constitutively express PPI modifiers (phosphatases) containing a chloroplast transit peptide, namely *cTP-SAC7.OX, cTP-PTEN.OX*, and *cTP-dOCRL.OX*. As previously mentioned, SAC7, PTEN, and dOCRL are capable of reversibly dephosphorylating PI4P, PI3P, and PI5P, respectively (Fig. **2a**). Of note, none of the transgenic lines expressing *cTP-SAC7, cTP-PTEN,* and *cTP-dOCRL* showed any visible developmental phenotypes, nor did they exhibit any abnormalities (Fig. **5b**). This observation aligns with the absence of binding of PPI biosensors to any PPIs in the chloroplast of plants growing under optimal conditions (Fig. **4a**). Subsequently, plant overexpressing plastidial phosphatase were subjected to drought stress by withholding water. As shown in Fig. **5b**, plants overexpressing *cTP-SAC7*, *cTP-PTEN*, and *cTP-dOCRL* began to exhibit leaf wilting symptoms approximately 8 days into the drought period. These symptoms became progressively more pronounced in the subsequent days of drought compared to the control plants. Furthermore, the measurements of the maximum quantum efficiency of photosystem (PS) II (Fv/Fm) in drought-treated plants revealed a photosynthetic impairment in plants expressing *cTP-SAC7*, *cTP-PTEN*, and *cTP-dOCRL* when compared to WT (Fig. **5b**). In addition, exposing seedlings overexpressing *cTP-PTEN* to 30°C for 9 days resulted in pale cotyledons as compared to WT (Supporting Information Fig. S5b). Next, we performed an ultrastructural analysis of plants overexpressing PPI modifying enzymes targeted to the plastids. No significant differences were observed compared to WT plants when grown under normal growth conditions (Fig. **5b, c** and Supporting Information Fig. S5c). In contrast, ultrastructural analysis of drought treated *cTP-PTEN* overexpression lines showed a strong swelling of the stromal fraction and thylakoid membranes compared to drought treated WT, indicating altered osmotic pressure within plastids of *PTEN.OX* plants (Fig. **5c**). Interestingly, similar phenotypes have been described for *VIPP1* knockdown lines (Zhang et al., 2012). Of note, we showed that VIPP1 and PI3P sensor protein colocalized (Fig. **3**).

**Figure 5:**
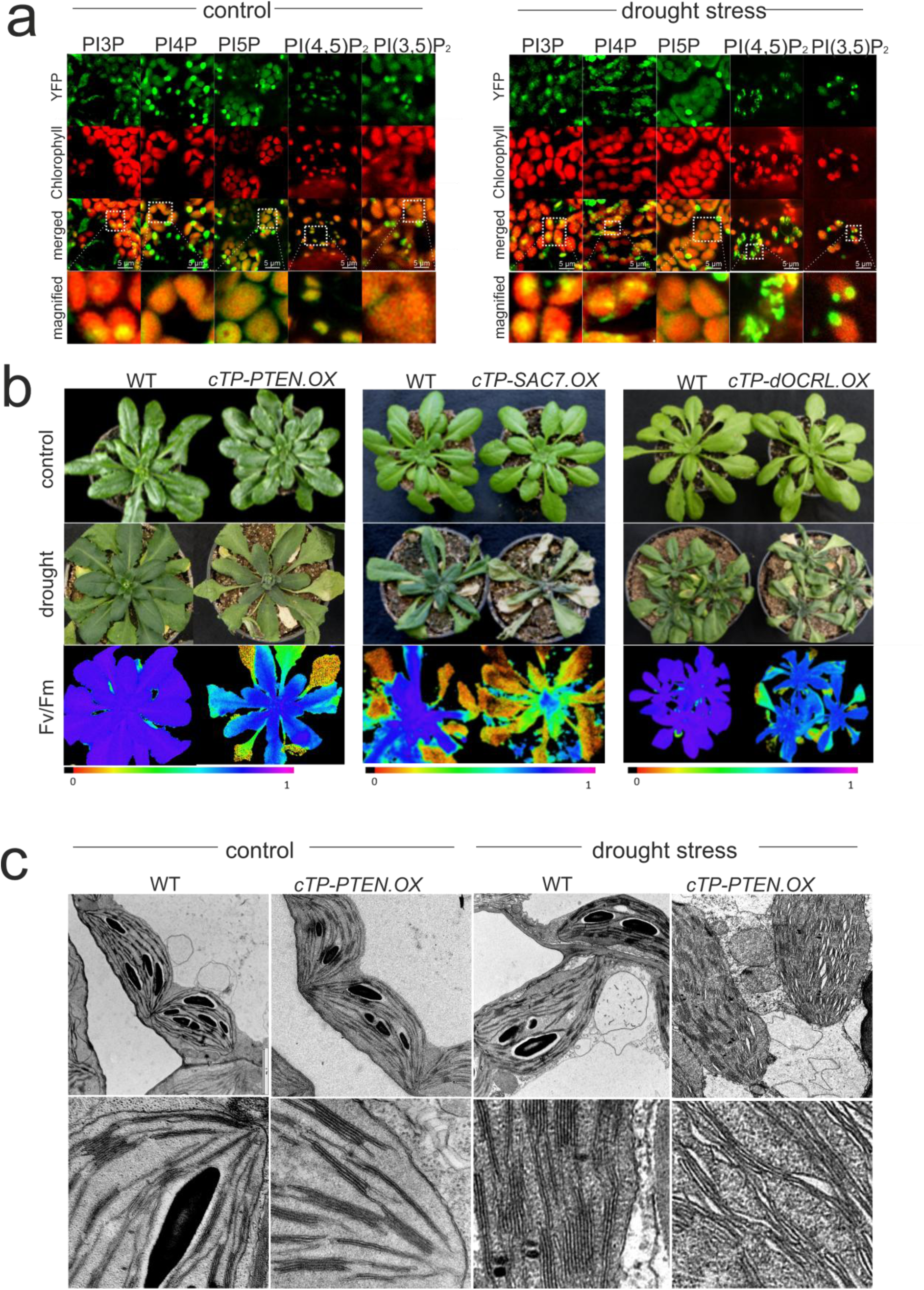
Perturbation of PIP abundance leads to drought sensitivity in *Arabidopsis* plants expressing PI4P phosphatase (SAC7), PI3P phosphatase (PTEN), and PI5P phosphatase (dOCRL). A. Sub-organelle distribution pattern of mEOS4b-PI3P, YFP-PI4P, YFP-PI5P, YFP-PI(4,5)P_2_ and YFP-PI(3,5)P_2_ biosensors in the chloroplast of *Arabidopsis* seedlings under control and drought stress conditions. Confocal images of 10-days-old *Arabidopsis* seedlings stably expressing *cTP-mEOS4b-PI3P*, *cTP-YFP-PI4P*, *cTP-YFP-PI5P*, *cTP-YFP-PI(4,5)P_2_* and c*TP-YFP-PI(3,5)P_2_* biosensors. For drought stress leaf samples from plants growing on petri dishes were detached and kept for 30 minutes at RT prior to imaging. B. Phenotypes of WT, *cTP-PTEN.OX* (left), *cTP-SAC7.OX* (middle) and *cTP-DOCRL.OX* (right) plants under control condition (well watered; upper panel) and after drought stress (middle panel). Seedlings were grown for 4 weeks in soil under a diurnal cycle of 16 h light (120 μE m^−2^ s^−1^) and 8 h dark at 22 °C. Drought stress was applied by withholding water for 10-14 days. The lower panel indicates *Fv/Fm* measurements of drought-stressed *cTP-PTEN.OX* (left) and *cTP-SAC7.OX* (right) plants. C. TEM images of *Arabidopsis* WT and *cTP-PTEN.OX* (left two rows). TEM images of Arabidopsis WT and plants expressing *cTP-PTEN.OX* following drought treatments (right two rows).

Taken together this data suggest that PPIs may serve as lipid signaling molecules within chloroplasts, fine-tuning the response to abiotic stresses.

## Discussion

Chloroplasts contain the most important membrane system in plant cells, thylakoid membranes, where the light-dependent reaction of photosynthesis takes place. Notably, chloroplast membranes show a distinctive lipid composition, mainly composed out of galactolipids and sulfolipids and very few phospholipids. Whereas recent studies have shown the bulk lipid composition of chloroplasts (LaBrant et al., 2018), presence and distribution of minor or signaling lipids remains elusive. Hence, a better understanding of the lipid composition of thylakoids and more broadly chloroplasts is currently crucial to understand their biogenesis maintenance driving photosynthesis under steady state and stress conditions. To date, presence of PPIs in the lipid composition of chloroplast membranes remains vague. This is most probably due to their low abundance and high turnover rates, which make them difficult to detect. Here, we show the detection of PPIs within the chloroplasts of tobacco (Fig. **1** and **2**) and heat stressed *Arabidopsis* plants (Fig. **4**), using our PPI biosensors *in vivo*.

Synthesis and turnover of PPI pools are vital for regulating various membrane trafficking processes membrane maintenance along with signaling transduction pathways (Meijer and Munnik, 2003; Di Paolo and De Camilli, 2006; Heilmann, 2009). Therefore, it is likely that PPI could play a similar role in chloroplasts. However, it remains unclear if PPIs can be produced within the chloroplasts. If so, it suggests the potential presence of unknown PI kinases and PPI phosphatases within this organelle. ARC3, a chloroplast division factor, represents a chimera of prokaryotic FtsZ and part of eukaryotic phosphatidyl-4-phosphate 5-kinase and localizes to the stroma (Shimada et al. 2004, Maple et al., 2007). Whether it functions as PI Kinase remains so far unexplored. Alternatively, PPI might be transferred from ER to chloroplast envelopes through membrane contact sites (LaBrant et al., 2018). However, the exchange of lipids between ER and plastid outer envelope membranes is highly restricted (Karkie et., 2019). Lipid exchange mediated by the action of flipases and flopases had been speculated (Rajasekharan and Gummadi, 2011). Once inside the inner leaflet of the inner envelope membrane PPIs could subsequently be transported within the chloroplast via the action of lipid binding proteins or envelope vesicles (LaBrant et al., 2018, Hertle et al., 2021, Schroda et al., 2021). So far, interactions between PPIs and chloroplast proteins, PDV1 and PDV2, Chloroplast Sec14-like protein 1 (CPSFL1), WHEAT KINASE START1 (WKS1), and VIPP1 have only been shown *in vitro*, using protein-lipid blot overlay assays (Gou et al., 2015; Okazaki et al., 2015; Theis et al., 2019; Hertle et al., 2020). PDV1 and PDV2 are integral components of the chloroplast division machinery, and they are responsible for regulating the rate of chloroplast division (Okazaki et al., 2009). CPSFL1 plays a crucial role in ensuring proper chloroplast development, promoting plant growth, and supporting optimal photosynthetic function in Arabidopsis and *Chlamydomonas* (García-Cerdán et al., 2020; Hertle et al., 2020; Kim et al., 2022). WKS1 provides resistance to *Puccinia striiformis f. sp tritici* in wheat (*Triticum aestivum).* VIPP1 plays essential roles in thylakoid membrane biogenesis, maintenance and repair under both normal and stress conditions in *Arabidopsis* and *Chlamydomonas* (Nordhues et al., 2012; Zhang et al., 2016; Theis et al., 2020). Here we demonstrated the co-localization of VIPP1 and the PI3P biosensor within the chloroplasts (Fig. **3a**). Additionally, we observed alterations in the sub-chloroplast distribution pattern of VIPP1 in the presence of the PI3P modifier, PTEN (Fig. **3b**). Our findings showed that plants expressing *cTP-PTEN* were sensitive to heat and drought stress and exhibited reduced photosynthetic efficiency (Fig. **5b**). Interestingly, ultrastructural analysis of *ctp-PTEN* overexpressing plants indicated osmolarity changes by swollen plastids and thylakoids (Fig. **5c**). Given the critical role of VIPP1 in thylakoid maintenance and repair under stress conditions (Zhang et al., 2016; Theis et al., 2020), it is possible that PI3P may influence the function of VIPP1 under stress conditions. However, we did not detect any changes in VIPP1 stability or its association with multiple complexes in *cTP-PTEN-*expressing plants (Supporting Information Fig. S3a, b). Nevertheless, the *in vivo* association of PI3P with VIPP1 (Fig. **3**) might fine-tune the function of VIPP1 under stress conditions.

In fact, phosphoinositide signaling pathways are a pivotal component mediating plant responses to various forms of stresses (Sui et al., 2008; Xue et al., 2009; Munnik and Vermeer, 2010; Zhai et al., 2013; Morales et al., 2019). Extensive research has confirmed that elevated temperatures trigger PPI flux in plant cells. This phenomenon is not limited to the model plant *Arabidopsis* (Mishkind et al., 2009; Annum et al., 2023) but extends across the plant kingdom, encompassing species such as rice (Mishkind et al., 2009), wheat (Annum et al., 2022)*, Cycas multipinnata*, and *C. panzhihuaensis* (Zhu et al., 2022). Hence, the inducibility of PPI flux within the cells in response to heat stress appears to be evolutionarily conserved, indicating the crucial role of these phospholipids in the response to high-temperature stress. We verified, that high temperature and subsequently induction of PPI flux within the cell leads to relocation of PPI biosensors, indicating an increase in PPI concentration inside the chloroplasts (Fig. **4**). Thus, changing cytoplasmic PI or PPI pools most probably also affects PI and PPI flux in the plastid.

Drought induces water deficiency and hinders plant growth. Numerous studies have illustrated the crucial role of the phosphoinositide signaling pathway in drought stress across different plants (Mikami et al., 1998; Perera et al., 2008; Wang et al., 2008; Zhai et al., 2013; Carpaneto et al., 2017). The instant induction of *AtPIP5K1*, Phosphatidylinositol-4-phosphate 5-kinase, transcript in Arabidopsis plants subjected to drought or abscisic acid treatment suggests the participation of AtPIP5K1 in altering PPI pools and, consequently, its involvement in water-stress signal transduction (Mikami et al., 1998). Furthermore, dehydration induces the levels of PI5P through the activity of a myotubularin homolog (*AtMTM1*). MTM1 is itself induced during dehydration and can produce PI5P from PI(3,5)P_2_ (Ndamukong et al., 2010). Moreover, it has been reported that drought stress induces synthesis of PI(3,5)P_2_ in plants (Meijer et al., 1999) in which PI(3,5)P_2_ controls the decrease in vacuolar volume to counteract water deficiency and facilitates the closure of stomata (Carpaneto et al., 2017). Analysis of expression patterns showed that six out of nine Arabidopsis *PLC* genes are induced by dehydration (Tasma et al., 2008). Similarly, the expression of maize *ZmPLC1* is induced under dehydration, and this increased expression contributes to the improved drought tolerance of maize (Wang et al., 2008). In addition, it has been reported that *Arabidopsis* transgenic plants, overexpressing mammalian inositol polyphosphate 5-phosphatase display modified responses to abscisic acid and significantly improved drought tolerance (Perera et al., 2008). These studies provide evidence for a close association between phosphoinositide signaling pathways and water-stress responses in plants. We observed that perturbation of PPI balance in chloroplasts upon dehydration in plant overexpressing different PPI phosphatases resulted in drought sensitivity (Fig. **5**). Our results suggest the generation and turnover of PPI pools within the chloroplasts may play crucial roles in regulating signaling transduction pathways.

The alterations in PPI levels or localization triggered by dehydration can be detected by downstream protein targets possessing PPI-binding domains, consequently altering their activity or localization (Munnik and Vermeer, 2010). For instance, the activity of the *Arabidopsis* trithorax-like factor ATX1 was modified upon dehydration stress due to the binding of PI5P (Ndamukong et al., 2010). Hence, further research is required to comprehensively characterize chloroplast proteins possessing PPI binding domains.

While organelles have distinct specialized functions as individual compartments, there is evident that inter-organelle communication plays a vital role in preserving cellular function. This collaborative interaction occurs through various membrane-mediated processes and signaling molecules. It has been reported that phospholipids are a notable class of molecules that promote these interactions (Meijer and Munnik, 2003; Ravi et al., 2021). Hence, it is reasonable to hypothesize that PPIs may play a role in mediating cross-communication between the chloroplast and other cellular organelles, such as the nucleus. Comprehensive research efforts are required to elucidate the involvement of PPIs in orchestrating signaling pathways and regulating crosstalk between chloroplasts and other organelles.

## Material and methods

### Generation of transgenic plants

For fluorescent reporters the cTP of RbcS (sequence in supplement) was amplified using oligos (AH1 and AH2) containing recombination sites for InFusion® cloning into pDK366 linearized by SpeI. Resulting plasmids were linearized by SpeI and synthetic DNA of YFP, mCherry, CFP or mEOS4b respectively (and codon optimized for expression in A. thaliana) was inserted in frame to the n-terminal cTP by InFusion cloning. The resulting plasmids were used to create chloroplast targeted fluorescent lipid sensor and lipid modifying proteins. In brief, the coding sequence of the lipid binding domains of the pleckstrin homology (PH) domain of human ARHGEF9 for PI5P binding, the PH domain of human rho guanidine nucleotide exchange factor 7 (ARHGEF7) for PI(3,5)P_2_ detection (Singh et al., 2021), the four phosphate adaptor protein 1 and 2 (PH) domain of rat FAPP1 for PI4P detection, the PH domain of mouse 1-phosphatidylinositol-4,5-bisphosphate diesterase delta (Plcd1) for PI(4,5)P_2_ detection and putative phosphatidylinositol 3-phosphate binding site encoded by the FYVE domain from human hepatocyte growth factor related tyrosine kinase substrate (HGS11) (Simon et al., 2014) was codon optimized for expression in Arabidopsis were codon optimized and synthesized with recombination sites (sequences in supplement) to be integrated into pDK366 containing cTP_YFP, cTP_mCherry or cTP_mEOS4b fusion proteins under the control of the 35S promotor by InFusion® cloning. mCherry-FKBP-PTEN was a gift from Gerry Hammond (Addgene plasmid # 116866; http://n2t.net/addgene:116866; RRID:Addgene_116866), SAC7/RHD4 (At3g51460) was amplified from Arabidopsis thaliana cDNA. dOCRL was used as codon optimized synthetic DNA of the sequence publishes in Doumane et al., 2021. VIPP1 was amplified from Arabidopsis cDNA using oligos (AH15/AH16). Subsequent VIPP1 lacking the stop codon was fused in frame to codon optimized CFP (sequence in supplement) by fusion PCR using oligos coding for recombination sites for infusion cloning into pDK488 linearized by SpeI. For stable expression, we introduced the constructs into *Arabidopsis* (Col-0) through a floral dip method utilizing Agrobacterium tumefaciens (strain GV3101) (Clough and Bent, 1998). Transient expression in *N. benthamiana* was conducted following the established procedure (Li et al., 2011).

### Plant materials, growth conditions, and the procedures for heat and drought treatments

*Arabidopsis thaliana* Columbia (Col-0) was used as the WT. The seeds were germinated on 0.5× Murashige and Skoog agar medium enriched with 1% (w/v) sucrose. The seedlings were cultivated following a 16-hour light cycle (120 μmol m^−2^ s^−1^) at 22 °C, and 8 h dark at 22 °C. To induce elevated PPI levels in the cells, 10-day-old Arabidopsis seedlings were subjected to a heat regime of 40 °C for 30 min. To assess heat stress tolerance, 7-day-old Arabidopsis seedlings were exposed to a heat regime of 45.5 °C for 45 min. To assess drought tolerance, the seedlings were cultivated for 4 weeks in soil under a diurnal cycle comprising 16 hours of light (120 μmol m^−2^ s^−1^) and 8 hours of darkness at 22 °C. Subsequently, to induce drought stress, watering was discontinued for a period of 10-14 days.

### Chloroplast or thylakoid Isolation

To isolate chloroplasts, leaf material from the various genotypes was initially ground in chloroplast isolation buffer. The buffer composition included 0.45 M sorbitol, 20 mM Tricine (pH 8.4), 10 mM EDTA, 10 mM NaHCO3, and 0.1% BSA. Chloroplasts were collected by centrifugation for 5 min at 3000g, 4°C from the collected material obtained by filtering it through two layers of Miracloth. The released chloroplasts were then carefully layered onto a Percoll gradient made with 5X resuspension buffer (0.3 M sorbitol, 20 mM Tricine (pH 8.4), 2.5 mM EDTA, and 5 mM MgCl2). Centrifugation was performed in a swing-out rotor at 4°C for 10 minutes at 3300g. Intact chloroplasts were collected from the lower green band and subsequently subjected to a wash using 1X resuspension buffer.

Thylakoid membrane preparation from leaves of 3-week-old WT and *PTEN.OX* plants was conducted in accordance with the method described in (Bassi et al., 1985). Leaf material was homogenized in an ice-cold buffer comprising 0.4 M sorbitol, 0.1 M Tricine/KOH (pH 7.8), and 1 mM PMSF. The resulting homogenate was filtered through a nylon mesh and then centrifuged at 4°C, 3000g for 10 minutes. Subsequently, chloroplasts were disrupted in a buffer consisting of 20 mM HEPES/KOH (pH 7.8) and 10 mM EDTA (pH 8.0). Thylakoids were collected by centrifugation at 12,000g for 10 minutes and resuspended in a buffer containing 50% glycerol, 10 mM HEPES/KOH (pH 7.5), and 1 mM EDTA (pH 8.0). Fractionation of spinach chloroplasts was performed as described in (Hertle et. al 2020). Envelope fractions were separated into inner and outer envelopes using sucrose gradients as discribed in (Block et al., 2018).

### Thermolysin treatment

Intact chloroplasts were treated with thermolysin based on Fitzpatrick et al., 2001. The chloroplast pellet, equivalent to 30 μg of chlorophyll was resuspended in import buffer (330-mM sorbitol and 50 mM HEPES–KOH, pH 8.0) and incubated with thermolysin (36 units mg^−1^; 0.05 units µg^−1^ chlorophyll) for 30 minutes on ice. Intact chloroplasts were re-isolated over a 40% v/v Percoll. chloroplasts were dissolved in 100ul of 4xSDS buffer.

### Bis(sulfosuccinimidyl)suberate (BS3) treatment

Isolated thylakoids from WT and *PTEN.OX* were washed 5 times with 20 mM HEPES/ KOH pH 7.5 to remove the EDTA. Crosslinking was performed by adding 2mM BS3 to a thylakoid suspension containing 150 µg/ml of chlorophyll, followed by incubation on ice for 20 minutes. The reaction was stopped with 150 mM TRIS HCL pH:7.5. The plates were dissolved in 5X SDS sample buffer for further analysis.

### Protein extraction and immunoblotting

Soluble proteins were extracted from Arabidopsis seedlings expressing PPI biosensors following the method outlined in (Sedaghatmehr et al., 2016). The protein concentration was determined using the Bradford protein assay. Subsequently, the proteins were separated by SDS-PAGE, and the PageRule Plus Prestained Protein Ladder (10–250 kDa; Thermo Scientific) served as the molecular weight marker.

For immunoblot analysis, the proteins were transferred onto either a Protan nitrocellulose or Polyvinylidene Difluoride membrane. Primary antibodies used included a rabbit anti-VIPP1 polyclonal antibody from Agrisera, rabbit anti-TOC159 from Agrisera, and mouse anti-green fluorescent protein (GFP) antibody from Clontech. Secondary antibodies, IRDye 800CW-conjugated goat anti-rabbit or anti-mouse IgG (H+L), were employed at a 1:10,000 dilution (LI-COR Biosciences). Band detection was performed using the Odyssey Infrared Imaging System (LI-COR Biosciences).

### GUV formation

To form GUVs, PVA assisted swelling was performed in PBS buffer according to Weinberger et al., 2013. PVA-coated substrate was prepared by spreading and drying of 100–300 *μ*L of PVA (5%) solution on a glass slide. Afterwards 10–20 *μ*L of plant lipid mixtures (2 mM final) of MGDG/DGDG/SQDG/PG/PI (52:26:6.5:9.5:1) (v/v) according to Hölzl et al., 2019 Labrant et al., 2018; Block et al,.1983), with or without 0.1% PI3P and with 0.01mol% DiI. dissolved in chloroform were spead on the dried PVA film and placed under vacuum for 30 min for solvent evaporation. A chamber was formed using vitrex and a coverglass filled with sucrose (200 mM) containing PBS buffer. Following 15 min. incubation, GUVs were transferred into an Eppendorf tube using a pipette until further use. For protein binding studies proteins were added in PBS containing glucose in equal osmolarity in a final concentration of 0.5 µM.

### Expression of recombinant his tagged VIPP1-mEOS4b and FYVE-mEOS4b

VIPP1 lacking the cTP and the stop codon was amplified and fused to mEOS4b by fusion PCR using the primers AH13 and AH14. mEOS-FYVE was amplified from plant plasmids described above using primers overlapping to pET28b. The PCR product was integrated into pET28b linearized by restriction using BamHI and EcoRI by in Fusion cloning. Resulting plasmid was used for inducible expression of 6xHis tagged VIPP1-mEOS. VIPP1 was expressed by adding 0,5 mM IPTG to a culture of O.D.600=0.5 grown at 37°C. The temperature was lowered to 30°C and the culture was grown for 3 more hours. The bacteria were harvested by centrifugation and the pellet was snap frozen in liquid nitrogen and stored at −70°C until further use. Recombinant protein purification was performed using Ni-NTA(Qiagen) according to the manufacturers protocol for native proteins. The lysis buffer was supplemented with 2% Triton-X-100 (Serva). To remove access detergent and imidazole, buffer exchange of the obtained purified protein solution was achieved by 10x centrifugation of the solution through a 10.000 MW cut off centricon (Millipore) using PBS. The protein was stored at −70°C until used.

### Fat blot assays

Fat blot assays were performed as described in (Munnik and Wierzchowiecka, 2013). FYVE-mEOS was detected using a gel doc fluorescence blot reader (Biorad) at 488 nm excitation.

### TEM microscopy

Leaf samples were fixed in 2.5% (w/v) glutaraldehyde, 2% (w/v) paraformaldehyde in 50 mM sodium cacodylate (pH 7.4) for 1 hr under vacuum at RT. Subsequent fixation was continued overnight at 4°C. After washing in PBS Post-fixation with was performed in 2% OsO4 and 0.8% K_3_Fe(CN)_6_ in 50 mM cacodylate buffer (pH 7.4) for 2 hr at RT. After rinsing, leaf samples were post-stained with 2% (w/v) uranyl acetate for 2 h at RT. Subsequent dehydration was performed using acetone and Spurr’s resin (Science Services) was infiltered. Polymerisation was carried out at 60°C for 48hrs. Ultrathin sections (50-70 nm) were cut on a UC-6 ultramicrotome (Leica) using a diamond knife. For transmission electron microscopy, sections were deposited on formvar and carbon coated nickel grids and contrasted with 2% <u>uranyl acetate</u> and Reynold’s lead citrate. Images were acquired on a JEOL JEM F-200 (JEOL).

### Confocal microscopy

Plants were imaged using a Leica TCS SP8 or SP5 confocal microscope. The excitation/emission settings used were 405/415-475nm for CFP, 488/505-535 nm for GFP, 512/520-540 nm for YFP and mEOS4b, 561/570-600 nm for RFP and DiI, and 633/650-720 nm for chlorophyll fluorescence. All images were acquired using sequential scanning settings. The images analysis and quantification of fluorescence intensities was performed using the ImageJ software.

## Acknowledgments

We thank the Deutsche Forschungsgemeinschaft (DFG) and the Max Planck Society (MPG) for funding. We thank Prof. Dr. Ralph Bock and his team from the Max Planck Institute of Molecular Plant Physiology (MPIMP, Potsdam, Germany) for supporting and hosting the project and engaging in insightful discussions.

We thank Dr. Nadja Tarakina and Bolortuya Badamdorj from the Max Planck Institute of colloids and interfaces for providing access and excellent technical assistance to the transmission electron microscope.

We thank Dr. Rumiana Dimova and Dr. Shreya Pramanik at Max Planck Institute of colloids and interfaces (MPIKGF, Potsdam, Germany) for help and discussions with plant GUV experiments

We thank Prof. Petra Bauer at the Institute of Botany, at the Heinrich Heine University (Düsseldorf, Germany) for hosting the project and discussions.

## Author contributions

AH conceived the study. MS and AH designed and performed the experiments; FR generated PIP biosensor constructs. MS and AH wrote the manuscript. All authors agreed to the submission of the manuscript.

## Supporting Information

**Figure S1.** Intensity plots of PPI biosensors in chloroplasts following transient expression in tobacco.

**Figure S2.** Colocalization of PPI biosensors with two different fluorescent proteins in the chloroplasts.

**Figure S3.** VIPP1 stability and/or its multiple complexes in *PTEN-expressing* plants and PI3P detection in chloroplasts.

**Figure S4.** Distribution pattern of YFP-PI4P, YFP-PI5P, YFP-PI(4,5)P_2_, and YFP-PI(3,5)P_2_ biosensors in the chloroplasts of Arabidopsis under heat stress conditions.

**Figure S5.** Distribution pattern of YFP-PI4P, YFP-PI5P, YFP-PI(4,5)P_2_, and YFP-PI(3,5)P_2_ biosensors in chloroplasts of Arabidopsis under drought stress conditions.

**Supplementary Table 1.** Oligos used in the study

**Supplementary Table 2.** Sequences of synthetic DNA or fluorescent reporters used in the study

