## Supplemental Information for "New evidence for the presence and function of phosphoinositides (PPIs) in the chloroplast"

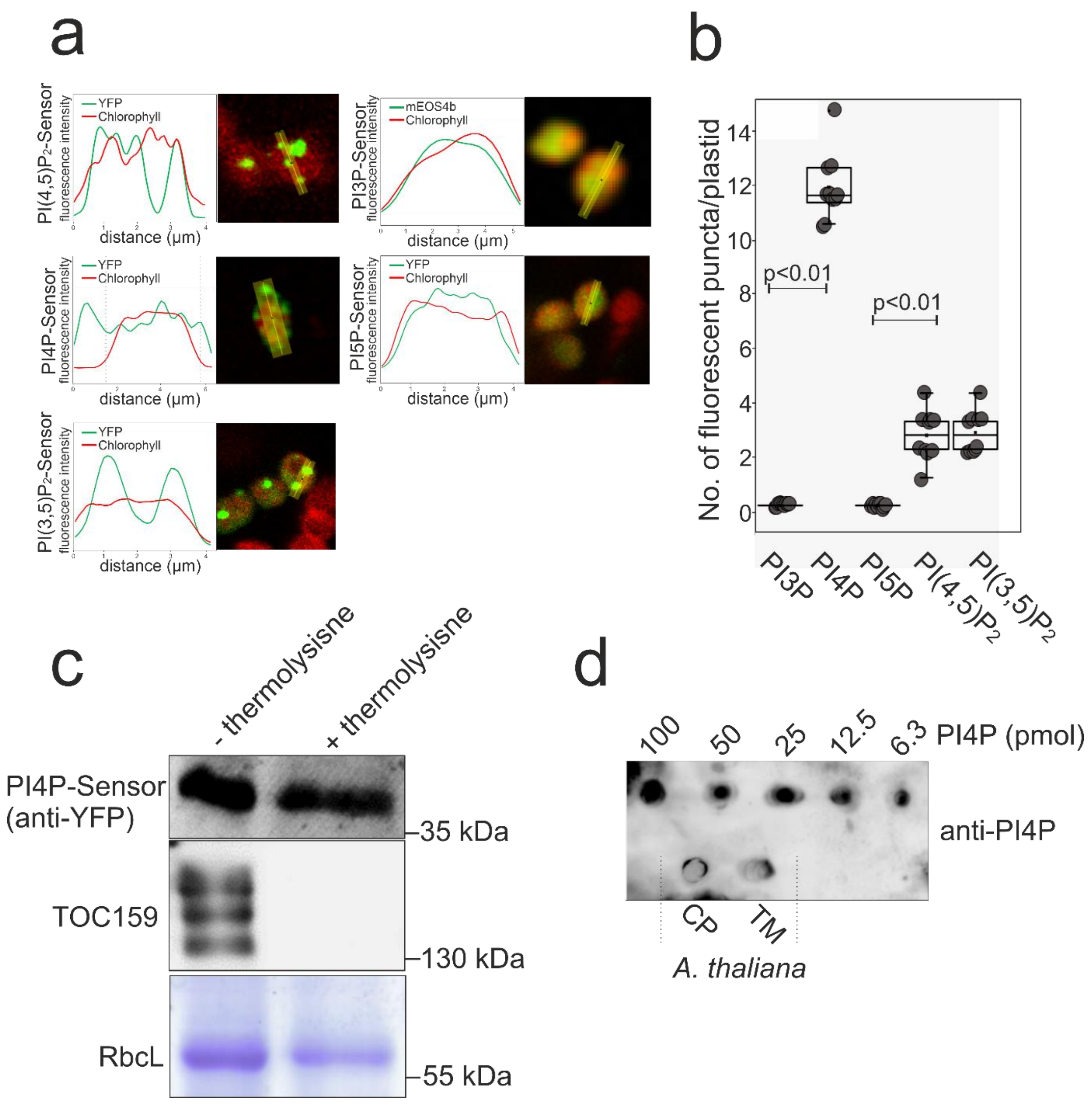


**Supplementary Fig. S1** a. Intensity blots of PPI biosensors in chloroplasts following transient expression in tobacco. Fluorescence intensity along the yellow line depicted in the microscopy image (right) is plotted together with chlorophyll fluorescence. b. experiments have been performed at least three times with identical results (n>3). The number of fluorescent puncta of 10 chloroplasts each sensor was counted and compared among the sensors using a paired t-test. P values below 0,01 are indicated (p<0,01). c. YFP-FAPP (PI4P) labeled structures are not localized at the outer envelope membrane. Intact chloroplasts of Arabidopsis plants expressing *YFP-FAPP* were subjected to thermolysin treatment (+) or control-treated (-) for 30 minutes. Proteins from chloroplasts were separated by SDS-PAGE followed by western blot against YFP and TOC159 antibodies. d. Fat blot assay on chloroplast and thylakoid lipid extracts together with PI4P lipid standard (ranging from 100 to 6.3 pmol) showed recognition of PI4P using PI4P antibodies.

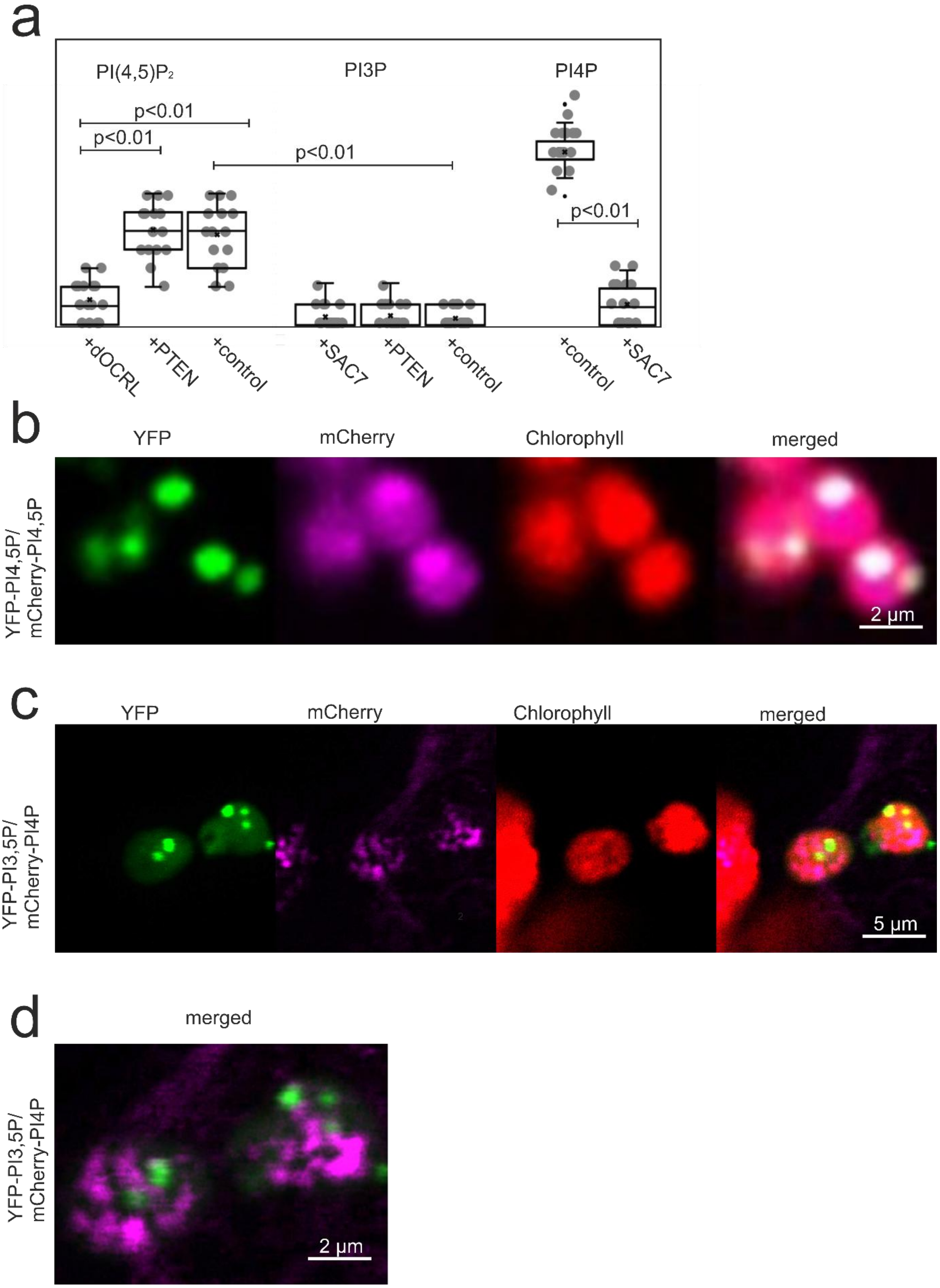


**Supplementary Fig. S2** Co-localization of PPI biosensors with two different fluorescent proteins in the chloroplasts. a. Quantification and statistical evaluation of PPI sensor fluorescence. Number of fluorescent puncta of 20 chloroplasts each was compared. Experiments have been performed more than three times (n>3). Statistical significance is indicated by paired t-test with a p value below 0.01($p<0.01$). b. Confocal images of tobacco leaves transiently expressing *cTP-YFP-PI(4,5)P*$_2$ and *cTP-mCh- PI(4,5)P*$_2$. c. Confocal images of tobacco leaves transiently expressing *cTP-YFP-PI(3,5)P*$_2$ and *cTP-mCh- PI4P*. d. Higher magnification of chloroplasts from B without chlorophyll shows no colocalization of the fluorescent proteins.

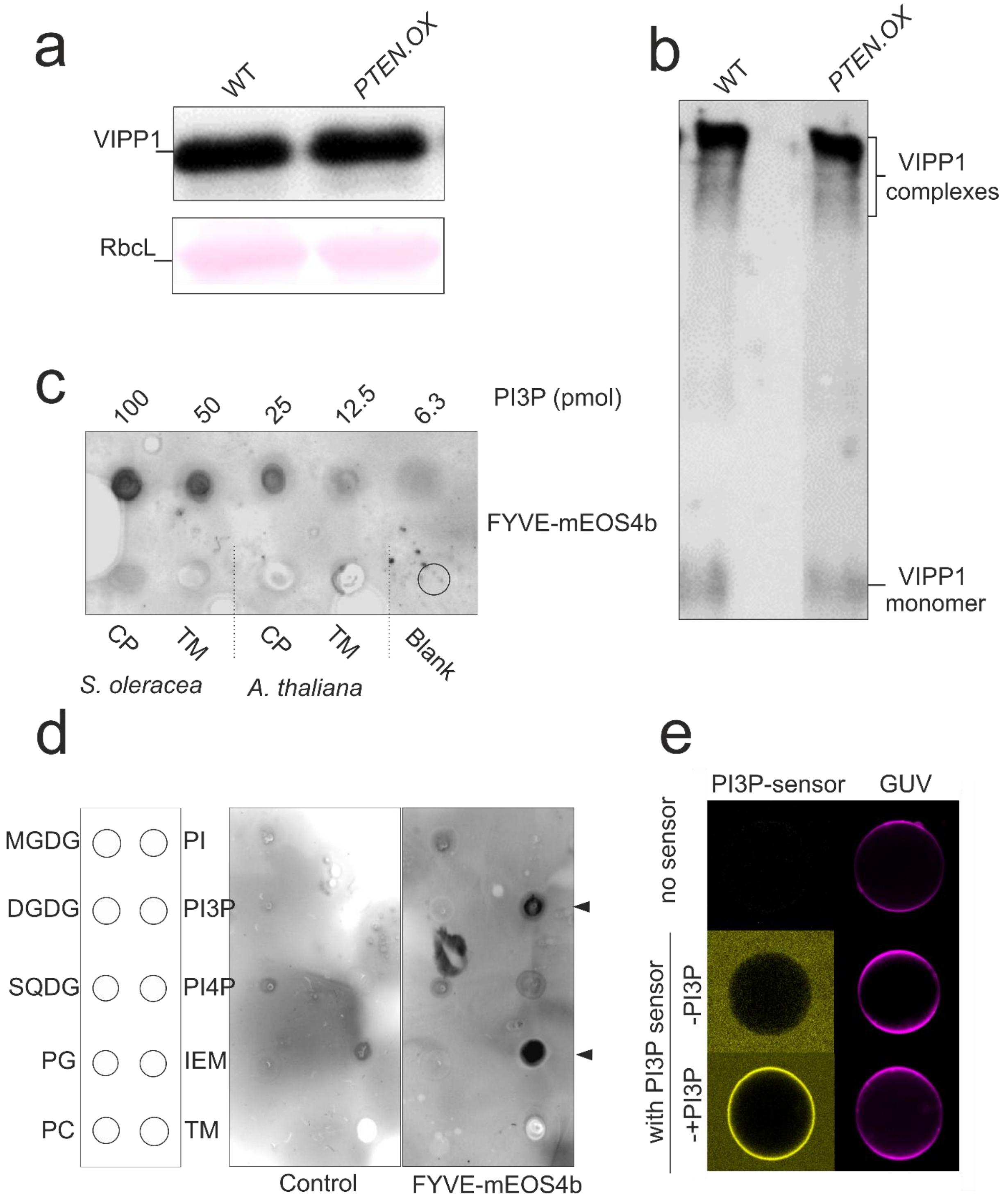

**Supplementary Fig. S3** a. VIPP1 stability and/or its multiple complexes in *cTP-PTEN-expressing* plants and PI3P detection in chloroplasts. Immunoblot analysis of VIPP1 in WT and *cTP-PTEN.OX* plants using VIPP1 antibodies. b. Thylakoid proteins from both WT and *PTEN.OX* plants were crosslinked with BS3, separated by SDS-PAGE, transferred to PVDF membranes, and subsequently immunolabeled using VIPP1 antibodies. c. Fat blot assay on lipid extracts of chloroplasts and thylakoid preparations from spinach and *Arabidopsis* using PI3P lipid standards and recombinant FYVE sensors (FYVE-mEOS4b). d. PI3P is specifically and exclusively recognized by recombinant FYVE sensors in the presence of chloroplast lipids (MGDG, DGDG, SQDG, PG, PC and PI). PI3P sensors strongly recognize inner envelope membranes and thylakoids of spinach chloroplast preparations. e. Recombinant FYVE-mEOS4b sensors bind to GUVs made from chloroplast lipids only in the presence of PI3P.

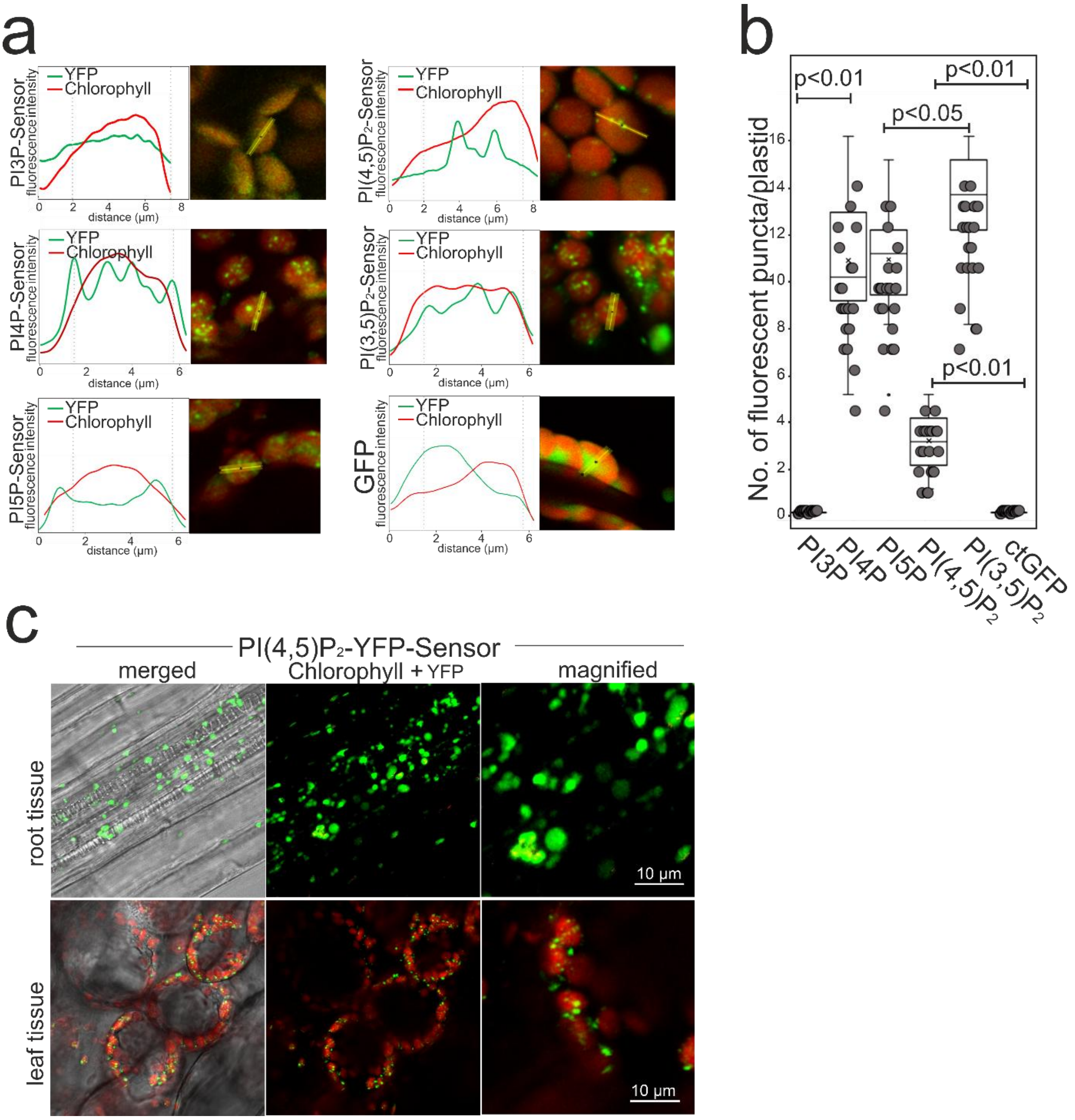

**Supplementary Fig. S4** Distribution pattern and intensity plots of YFP-PI4P, YFP-PI5P, YFP-PI(4,5)$P_2$ and YFP-PI(3,5)$P_2$ biosensors in the chloroplasts of *Arabidopsis* under heat stress conditions (at 40°C for 30 minutes). Confocal images of 10-days- old Arabidopsis leaves stably expressing *cTP-YFP-PI4P*, *cTP-YFP-PI5P*, *cTP-YFP-PI(4,5)P$_2$* and c*TP-YFP-PI(3,5)P$_2$* biosensors. a. Biosensor fluorescence along the yellow line depicted in the microscopy image (right) was plotted together with chlorophyll fluorescence. b. For each biosensor line the number of fluorescent puncta per chloroplast (n=20) was counted and compared statistically using a paired t-test. Significant differences are indicated when p values are below 0,01 ($p<0.01$). c. Distribution pattern of YFP-PI(4,5)$P_2$ biosensors in chloroplasts and the plastids of *Arabidopsis* roots under control and heat stress (HS) conditions. Confocal images of 10-days-old Arabidopsis leaves and roots stably expressing *cTP-YFP-PI(4,5)P$_2$* biosensors. Only in chloroplasts PI(4,5)P2 sensors form punctate structures.

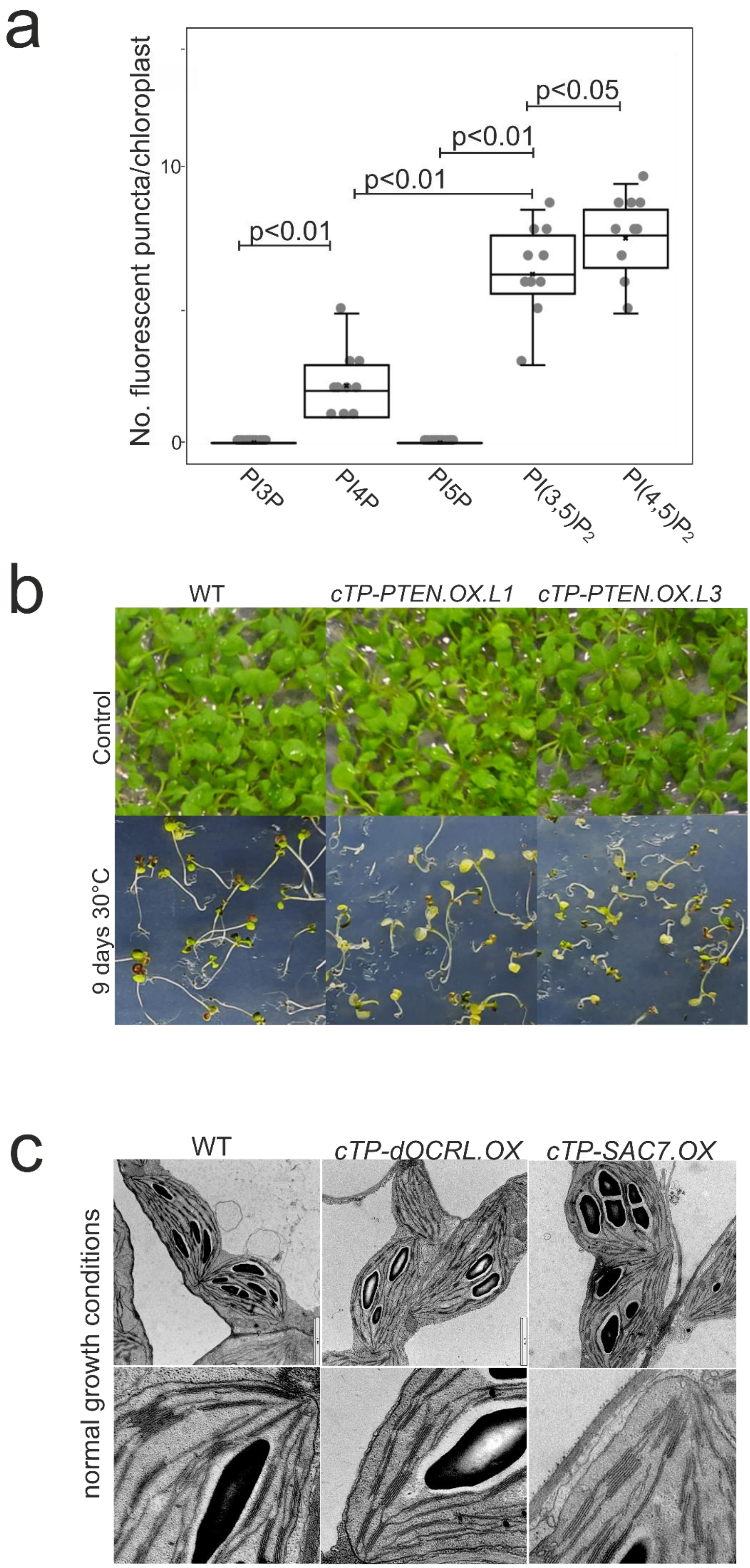
a
No. fluorescent puncta/chloroplast
0
10
p<0.01
p<0.01
p<0.01
p<0.05
PI3P
PI4P
PI5P
PI(3,5)P2
PI(4,5)P2
b
WT
cTP-PTEN.OX.L1
cTP-PTEN.OX.L3
Control
9 days 30°C
c
WT
cTP-dOCRL.OX
cTP-SAC7.OX
normal growth conditions

**Supplementary Fig. S5** a. Phenotype of WT seedlings and two independent PI3P phosphatase overexpressing lines (*cTP-PTEN.OX.L1* and *cTP-PTEN.OX.L3*) grown under normal conditions and under heat stress (9 days at 30°C). b. Quantification of fluorescent puncta in chloroplasts formed by PPI biosensors was analyzed for each biosensor line (n chloroplast=20). Numbers were compared statistically using a paired t-test. Significant differences are indicated when p values are below 0,01 ($p<0.01$). c. Electron micrographs of chloroplasts from WT and chloroplast PPI phosphatase overexpressing plants. *cTP-dOCRL.OX* expresses a chloroplast targeted $PI(4,5)P_2$ phosphatase. *cTP-SAC7.OX* expresses a chloroplast targeted PI4P phosphatase.

| | | |
|---|---|---|
| AH1 | cTP_F | 5´-gattaacactgaattcatggcttcctctatgctctc-3´ |
| AH2 | cTP_R | 5´-tctggtttgatagcagaaactggccacaccttcatgcagc-3´ |
| AH3 | mEOS4b_F | 5´-gctgcatgaaggtgtggccagtttctgctatcaaaccaga-3´ |
| AH4 | mEOS4b_R | 5´-cagggatatcactagtgctgccaccagatcttctagcgttatcaggaag-3´ |
| AH5 | mCherry_F | 5´-atggtttctaaaggtaaagtgagcaaggcgaggagga-3´ |
| AH6 | mCHerry_R | 5´-cagggatatcactagtcttgtacagctcgtccatgc-3´ |
| AH7 | YFP_F | 5´-gcatggacgagctgtacaagatggagacggttgatt-3´ |
| AH8 | YFP_R | 5´-cgcccttgctcaccatgttaggccaaaccttcatgcaag-3´ |
| AH9 | CFP_F | 5´-catcatcatcaccatcatgtgag-3´ |
| AH10 | CFP_R | 5´-cagggatatcactagtttacttgtaaagttcgtccattcc-3´ |
| AH11 | Sensor_F | 5´-gcatggacgagctgtacaagggcagcactagtaaaggatc-3´ |
| AH12 | Sensor_R | 5´-ctcgaattcgcggccgcttagctagtgatatccctgtg-3´ |
| AH13 | pET_VIPP1-mEOS_ | 5´-aaatgggtcgcggatccgaa atgaatctttttgaacgatt-3´ |
| AH14 | pET_VIPP1-mEOS_ | 5´-ttgtcgacggagctcgaattcttctagcgttatcaggaag-3´ |
| AH15 | VIPP1_F | 5´-gattaacactgaattcatggctctcaaagcttcacctg-3´ |
| AH16 | VIPP1_R | 5´-atggtgatgatgatgaaagtcgttagctttccttcgca-3´ |

**Supplementary Table I**

Oligos used in the study

cTP:

ATGGCTTCCTCTATGCTCTCCTCTGCCGCTGTGGTTACCTCCCCGGCTCAAGCCACCAT
GGTCGCTCCATTACTGGTTTGAAGTCATCCGCTTCTTTCCCGGTCACCCGCAAGGCCAA
CAACGACATTACTTCCATCACAAGCAATGGGGGAAGAGTTAGCTGCATAAGGTGTGGCC
A

mEOS4a:

ATGGTTTCTGCTATCAAACCAGATATGAGAATCAAGCTTAGAATGGAGGGAAATGTTAAT
GGACATCATTTTGTTATCGATGGAGATGGAACTGGAAAACCATATGAGGGTAAGCAAAC
TATGGATCTTGAAGTTAAAGAAGGAGGTCCTTTGCCATTCGCTTTTGATATTCTTACTACT
GCTTTCCATTACGGTAACAGAGTTTTTGTTAAGTATCCTGATAATATTCAAGATTATTTCA
AACAATCTTTCCCAAAAGGATACTCTTGGGAGAGATCTCTTACATTCGAAGATGGAGGAA

TTTGTTACGCTAGAAACGATATCACTATGGAGGGAGATACATTTTACAATAAAGTTAGAT
TCTATGGTACAAATTTCCCAGCTAATGGTCCAGTTATGCAAAAGAAAACTTTGAAATGGG
AGCCTTCTACAGAGAAGATGTACGTTAGAGATGGAGTTTTGACAGGTGATATCCATATG
GCTCTTCTTTTGGAAGGAAATGCTCATTACAGATGTGATTTCAGAACTACATATAAAGCT
AAAGAGAAGGGTGTTAAATTGCCAGGATACCATTTCGTTGATCATGCTATTGAGATCTTG
TCTCATGATAAGGATTACAACAAAGTTAAACTTTATGAACATGCTGTTGCTCATTCTGGA
CTTCCTGATAACGCTAGAAGA

YFP:

ATGGTGAGCAAGGGCGAGGAGCTGTTCACCGGGGTGGTGCCCATCCTGGTCGAGCTG
GACGGCGACGTAAACGGCCACAAGTTCAGCGTGTCCGGCGAGGGCGAGGGCGATGCC
ACCTACGGCAAGCTGACCCTGAAGTTCATCTGCACCACCGGCAAGCTGCCCGTGCCCT
GGCCCACCCTCGTGACCACCTTCGGCTACGGCCTGCAGTGCTTCGCCCGCTACCCCGA
CCACATGAAGCAGCACGACTTCTTCAAGTCCGCCATGCCCGAAGGCTACGTCCAGGAG
CGCACCATCTTCTTCAAGGACGACGGCAACTACAAGACCCGCGCCGAGGTGAAGTTCG
AGGGCGACACCCTGGTGAACCGCATCGAGCTGAAGGGCATCGACTTCAAGGAGGACG
GCAACATCCTGGGGCACAAGCTGGAGTACAACTACAACAGCCACAACGTCTATATCATG
GCCGACAAGCAGAAGAACGGCATCAAGGTGAACTTCAAGATCCGCCACAACATCGAGG
ACGGCAGCGTGCAGCTCGCCGACCACTACCAGCAGAACACCCCCATCGGCGACGGCC
CCGTGCTGCTGCCCGACAACCACTACCTGAGCTACCAGTCCGCCCTGAGCAAAGACCC
CAACGAGAAGCGCGATCACATGGTCCTGCTGGAGTTCGTGACCGCCGCCGGGATCACT
CTCGGCATGGACGAGCTGTACAAGTAA

mCherry:

GTGAGCAAGGGCGAGGAGGATAACATGGCCATCATCAAGGAGTTCATGCGCTTCAAGG
TGCACATGGAGGGCTCCGTGAACGGCCACGAGTTCGAGATCGAGGGCGAGGGCGAGG
GACGTCCCTACGAGGGCACCCAGACCGCCAAGCTGAAGGTGACCAAGGGTGGCCCCC
TGCCCTTCGCCTGGGACATCCTGTCCCCTCAGTTCATGTACGGCTCCAAGGCCTACGT
GAAGCACCCCGCCGACATCCCCGACTACTTGAAGCTGTCCTTCCCCGAGGGCTTCAAG
TGGGAGCGCGTGATGAACTTCGAGGACGGCGGCGTGGTGACCGTGACCCAGGACTCC
TCCCTGCAGGACGGCGAGTTCATCTACAAGGTGAAGCTGCGCGGCACCAACTTCCCCT
CCGACGGCCCCGTAATGCAGAAGAAGACCATGGGCTGGGAGGCCTCCTCCGAGCGGA
TGTACCCCGAGGACGGCGCCCTGAAGGGCGAGATCAAGCAGAGGCTGAAGCTGAAGG
ACGGCGGCCACTACGACGCTGAGGTCAAGACCACCTACAAGGCCAAGAAGCCCGTGC
AGCTGCCCGGAGCCTACAACGTCAACATCAAGTTGGACATCACCTCCCACAACGAGGA
CTACACCATCGTGGAACAGTACGAACGCGCCGAGGGCCGCCACTCTACCGGCGGCAT
GGACGAGCTGTACAAG

CFP:

ATGCATCACCATCACCACCACGTGAGCAAGGGTGAAGAACTCTTCACTGGTGTTGTGCC
TATCCTCGTTGAGCTTGATGGTGACGTGAACGGGCATAAGTTCTCCGTTTCTGGTGAAG
GTGAGGGTGACGCTACTTACGGAAAGCTCACTCTCAAGTTCATCTGTACCACCGGTAAG
CTCCCTGTTCCTTGGCCTACTCTTGTGACTACTCTCTCTTGGGGTGTGCAATGCTTCGCT
AGATACCCTGATCACATGAAGCAGCACGATTTCTTCAAGAGCGCTATGCCTGAGGGATA
CGTGCAAGAGAGAACCATCTTCTTCAAGGACGACGGGAACTACAAGACTCGTGCTGAG
GTTAAGTTCGAAGGTGACACCCTCGTGAACAGGATCGAGCTTAAGGGCATCGACTTCAA
AGAGGACGGAAACATCCTCGGACACAAGCTCGAGTACAACGCTATCCATGGGAACGTG
TACATCACCGCTGACAAGCAGAAGAACGGCATCAAGGCTAACTTCGGGCTCAACTGCAA
CATCGAGGATGGATCTGTTCAGCTCGCTGATCACTACCAGCAGAACACACCTATTGGAG
ATGGACCTGTGCTTCTCCCTGACAACCACTACCTTTCTACCCAGAGCAAGCTCAGCAAG

GACCCTAATGAGAAGAGGGACCACATGGTGCTTCTCGAGTTTGTTACTGCTGCTGGAAT
CACCCTCGGAATGGACGAACTTTACAAGTAA

PI3P-LBD:

actagtaaaggatccaATGCCATGTTCGCTGCTGAAAGAGCCCCTGACTGGGTGGATGCTGAG
GAATGCCATCGGTGCAGAGTACAGTTTGGGGTGGTGACCCGCAAGCATCACTGCCGAG
CATGTGGGCAGATCTTCTGTGGCAAGTGCTCCTCCAAGTACTCCACCATCCCCAAGTTC
GGCATTGAGAAGGAGGTGCGCGTGTGTGAGCCCTGCTATGAGCAGCTGAACAAGAAGG
CATAGctagtgatatccctgtgtgaaattg

PI4P-LBD:

actagtaaaggatccgGGGTGTTGTACAAGTGGACCAACTATCTCACAGGCTGGCAGCCTCGT
TGGTTTGTTTTAGATAATGGAATCTTATCCTACTATGATTCACAAGATGATGTTTGCAAAG
GGAGCAAAGGAAGCATAAAGATGGCAGTTTGTGAAATTAAAGTTCATTCAGCAGACAAC
ACAAGAATGGAATTAATCATTCCTGGAGAGCAGCATTTCTACATGAAGGCAGTGAATGC
AGCTGAAAGACAGAGGTGGCTGGTCGCTCTGGGGAGCTCCAAAGCATGTTTGACTGAT
ACAAGGACTAAATAGctagtgatatccctgtgtgaaattg

PI5P-LBD:

tcggcatggacgagctgtacaagCTCGAGAACATCGACAAGATCGCTCAATGGCAGGCTTCTGTT
CTTGACTGGGAAGGTGAGGATATCTTGGACCGTTCCAGCGAGCTTATCTACACTGGTGA
AATGGCCTGGATCTACCAGCCGTATGGAAGAAACCAGCAGAGGGTGTTCTTCCTGTTCG
ATCATCAGATGGTGCTCTGCAAGAAGGACCTCATCAGAAGGGACATCCTCTACTACAAG
GGGCGTATCGACATGGACAAGTACGAGGTTGTGGATATCGAGGATGGAAGGGACGACG
ACTTCAACGTGTCAATGAAGAACGCCTTCAAGCTCCACAACAAAGAGACTGAAGAGATC
CACCTGTTCTTCGCCAAGAAGCTCGAAGAGAAGATCAGATGGCTCAGGGCCTTCAGGG
AAGAACGTAAGATGGTGCAAGAGGACTGAagcggccgcgaattcgag

PI(3,5)$P_2$-LBD:

tcggcatggacgagctgtacaagGTGACCTACATGTCTCAGGTTCTCATCCAGTGCGCTGGGTCT
GAAGAAAAGAACGAGAGATACCTCCTTCTCTTCCCGAACGTGCTCCTTATGCTTTCTGCT
AGCCCTAGGATGAGCGGCTTTATCTACCAAGGTAAGCTCCCTACTACCGGGATGACTAT
CACTAAGCTCGAGGACTCTGAGAACCACCGTAACGCTTTCGAGATCAGCGGATCTATGA
TCGAGCGTATCCTCGTGTCTTGCAACAACCAGCAGGATCTCCAAGAGTGGGTTGAGCAT
CTCCAAAAGCAGACCAAGTGA agcggccgcgaattcgag

PI(4,5)$P_2$-LBD:

actagtaaaggatcctCGGGCCGGGACTTCCTGACCCTGCACGGGCTCCAGGATGACCCGGA
CCTTCAGGCCCTTCTGAAGGGCAGCCAGCTTCTGAAGGTGAAGTCCAGCTCGTGGCGT
AGGGAACGCTTCTACAAGCTACAGGAGGACTGCAAGACCATCTGGCAGGAATCTCGAA
AGGTCATGAGGTCCCCGGAGTCGCAGCTGTTCTCCATCGAGGACATTCAGGAGGTACG
GATGGGACACCGCACAGAAGGCCTGGAGAAGTTTGCCCGAGACATCCCCGAGGATCG
ATGCTTCTCCATTGTCTTCAAGGACCAGCGCAACACCCTAGACCTCATTGCCCCATCAC
CAGCTGACGCTCAGCACTGGGTGCAGGGCCTGCGCAAGATCATCCACCACTCCGGCTC
CATGGACCAGCGGCAGAAGCTGCAGCACTGGATTCACTCCTGCTTGCGAAAGGCTGAT

AAAAACAAGGACAACAAGATGAACTTCAAGGAGCTGAAGGACTTCCTGAAGGAGCTCAA
CATCCAGTAGctagtgatatccctgtgtgaaattg

PTEN:

tcggcatggacgagctgtacaagCCATGGTGGCTCCTTTTACCGGGCTCAAGTCATCTGCTAGCTT
CCCTGTTACCAGAAAGGCCAACAACGACATCACCAGCATCACCTCTAACGGTGGAAGA
GTGTCTTGCATGAAGGTTTGGCCTCTCGTGTCTAAGGGTGAAGAGGACAACATGGCCAT
CATCAAAGAATTCATGAGGTTCAAGGTGCACATGGAAGGCTCTGTGAACGGACATGAGT
TCGAGATCGAAGGTGAAGGTGAGGGTAGACCTTACGAGGGAACTCAGACTGCTAAGCT
CAAGGTGACAAAAGGTGGACCTCTTCCGTTCGCTTGGGATATCCTTTCACCGCAGTTCA
TGTACGGCAGCAAGGCTTATGTTAAGCACCCTGCTGATATCCCGGACTACCTCAAGTTG
TCTTTCCCAGAGGGATTCAAGTGGGAGCGTGTGATGAATTTCGAGGATGGTGGTGTGG
TGACTGTGACCCAGGATTCTTCACTTCAGGACGGGGAGTTCATCTACAAGGTGAAGCTC
AGGGGAACGAACTTCCCTTCTGATGGACCTGTGATGCAGAAAAAGACGATGGGCTGGG
AAGCTTCGTCTGAGAGAATGTATCCTGAGGACGGTGCTCTCAAGGGTGAGATCAAGCA
GAGACTCAAGCTTAAGGACGGTGGACACTACGACGCTGAGGTTAAGACTACCTACAAG
GCTAAGAAGCCTGTTCAGCTCCCTGGGGCTTACAACGTTAACATCAAGCTGGACATCAC
CTCGCACAACGAGGATTACACTATCGTCGAGCAGTACGAGAGAGCTGAGGGTCGTCAT
TCTACTGGTGGAATGGACGAGCTTTACAAGAGCGGACTCAGATCTAGAAGTGCTGCTGC
TGGCGCTGGTGGTGCTGCTAGAGCTGCTCTTGGAGTTCAAGTCGAGACTATCTCTCCTG
GTGACGGAAGGACTTTTCCTAAGAGGGGTCAGACTTGCGTGGTGCACTATACTGGAAT
GCTTGAGGACGGGAAGAAGTTCGACTCTTCTAGGGATCGTAACAAGCCGTTCAAGTTCA
TGCTCGGGAAGCAAGAGGTTATCCGTGGATGGGAAGAGGGTGTTGCTCAGATGTCTGT
TGGACAGAGAGCCAAGCTTACCATCTCTCCGGATTATGCTTACGGTGCTACTGGACACC
CTGGAATCATTCCTCCACATGCTACTCTCGTGTTCGACGTGGAACTTCTCAAACTCGAAT
CTGCTGGTGGATCTGCCGGTGGTAGTGCTGGTGGTTCAGCTGGCGGATCAGCTGGTG
GACCTAGAGCACAAGCTTCTAACTCTGCTGCTATGACCGCGATTATCAAAGAGATCGTC
AGCCGTAACAAGAGGCGTTACCAAGAGGATGGATTCGACCTCGATCTCACCTACATCTA
CCCGAATATTATCGCCATGGGATTCCCTGCTGAGAGACTCGAAGGGGTGTACCGTAACA
ACATCGATGATGTGGTGCGTTTCCTCGACAGCAAGCACAAGAACCACTACAAGATCTAC
AACCTCTGCGCCGAGAGGCACTACGATACAGCTAAGTTCAACTGCCGTGTGGCTCAGT
ACCCATTCGAGGATCATAACCCACCTCAGCTCGAGTTGATCAAGCCTTTCTGCGAGGAT
CTTGACCAGTGGCTTTCTGAGGATGACAACCATGTTGCTGCCATCCATTGCAAGGCTGG
AAAGGGAAGAACTGGGGTGATGATCTGCGCTTACCTTCTCCACAGAGGCAAGTTCCTTA
AGGCTCAAGAGGCTCTCGACTTCTACGGTGAGGTGAGAACCAGAGACAAGAAGGGTGT
CACTATCCCATCTCAAAGGCGTTACGTGTACTACTACAGCTACCTCCTGAAGAACCACC
TCGATTACAGACCTGTGGCTCTCCTGTTCCACAAGATGATGTTCGAGACAATCCCGATG
TTCAGCGGAGGAACTTGTAACCCTCAGTTCGTTGTTTGCCAGCTTAAGGTGAAGATCTA
CTCCAGCAACAGCGGACCTACTAGGCGTGAGGACAAGTTTATGTACTTCGAGTTCCCAC
AGCCTCTTCCGGTTTGCGGAGATATCAAGGTTGAGTTCTTCCACAAGCAGAACAAGATG
CTCAAGAAAGACAAGATGTTCCACTTCTGGGTGAACACCTTCTTCATCCCTGGACCTGA
AGAGACTTCTGAGAAGGTCGAGAACGGATCTCTCTGCGATCAAGAGATCGACTCCATCT
GCTCTATCGAGAGGGCTGACAACGACAAAGAGTACCTCGTTTTGACCCTCACCAAGAAC
GATCTCGACAAGGCGAACAAGGATAAGGCCAACCGTTACTTCAGCCCGAACTTCAAGGT
CAAGCTCTACTTCACCAAGACCGTCGAGGAACCTTCTAACCCAGAGGCTTCTTCTTCTA
CCTCTGTGACCCCTGATGTGAGCGATAACGAGCCTGATCACTACAGGTACTCTGACACC
ACCGATTCTGATCCTGAGAACGAGCCTTTCGATGAGGATCAGCACACCCAGATCACTAA
GGTGTGAACTAGTagcggccgcgaattcgag

SAC7:

tcggcatggacgagctgtacaagATGGAGACGGTTGATTCTCGGAACAAATTGCATTCCCGGTTAC
GTTTATGGGAATTCCCCGATCAGTACATCATCGAGCCAGCTGATGGCTCAGGCTCTTCA
TGTTTGGATATTAGTCGTGTTGATGCTTCCATGAAGCTCATCGATCAAGTCCCAGAAAGC
AATTCCGTGCGTGTCCCTAAAATCCGTTCCATTTTTGGTGTGGTTGGGATGTTGAAGCTC
CTTGCAGGATCATACTTGGTGGTTGTAACTGAGAGTGAACGTGTTGGATCGTTTCTGGG
GCATCCTATATTCAAAGTTACAACGCTTAAGGTTCTTCCTTGTGATCATTCACTTAAAAAC
TCGCCTGAAGAACAGAAAAAGATGGAGACCGAGTTCTCCAAGCTCTTAAGTGTAGCAGA
AAAGACAACTGGTCTCTATTTCTCATATGAAGTTAACTTAACATTGAGCTTCCTCTGTGTT
GGAAGTTCTCAAGGGAGCCATTGGAAGAGATATTGTTGACATTACTCTGATTGCTAGGA
GATGCACTAGGAGAAACGGTACACGTATGTGGAGAAGAGGAGCGGATCTTGATGGCTA
TGTTGCTAATTTTGTGGAGACTGAGCAAATTGTACAGATGAATGGGTATACGTCATCATT
TGTTCAGGTCAGAGGATCCATGCCTTTTATGTGGGAGCAGGTTGTAGATCTGACTTACA
AGCCCAAGTTTGAAATTGTCCAACCTGAAGAAGCTAAACGGATAGCTGAGCGTCACTTT
TTGGACCTTAGGAAAAAATATGGATCAGTTTTGGCAGTTGATCTCGTTAACAAGCAAGGC
GGTGAGGGGCGCTTATGCGAGAAGTATGCTACTGTTATGCAACACATTACTGGGGATGA
TATAAGGTACCTGCACTTTGATTTCCATCAAATTTGTGGGCATATTCATTTTGAGCGCTTA
TCAATTCTGTACGAGCAGATCGAGGGTTTTCTCGAAAAGAATGGGTACTTTCTGTTGAAT
GAAAAGGGTGAGAAAATGAAGGAGCAGCTTGGTGTTGTCCGATCAAACTGCATAGATTG
CTTAGACCGTACAAATGTTACCCAGAGCATGATAGGTCGAAAGATGTTAGAAGTTCAACT
CAAAAGGATTGGAGTTTTTGGCGCTGAAGAAACCATAAGCTCACATCTGAATTTTGACGA
ACACTATAAAATATTATGGGCTAATCATGGTGATGAAATCAGCATTCAGTACTCTGGTAC
TCCTGCACTTAAAGGAGATTTCGTTAGGTATGGGCATCGGACTGCCCATGGTGTCCTTA
AAGATGGTTGGAGCTCCCTCAGACGCTATTACCTGAATAACTTTGCTGATGGAACTAAG
CAGGATGCAATTGATCTCTTGCAAGGGCACTACATAGTTGCTGTGAGCCGAGACATGGC
TCCTGTACCTCAAAAGGGAGGCCTTGAGGCTGTAGCCAACTTCCCAGTGGCATTGTTCG
TGGTTCTGATGAGTTTCTGGTTTGCAACAATGTCTCTAAAACAAACTGGGAGTGATTATA
AGCATAAGCACTTGTTCTTCTCATTGTTGTGGACGGGAATTTGCGTGGGCATGGCTGCA
CTGGTGAGGGCCAATGGCAGGATCTTCTGTAACCGACCTCGTCTGCACAAGCCCAGAG
GCTGAagcggccgcgaattcgag

dOCRL:

tcggcatggacgagctgtacaagGTAAAGCAAGAGTTAAAGAAGCGAGAGAGTGAGTATATCGTCT
ATAAGGACATTATCATTTACTGCGCTACGTGGAACGTGAACAATAAGACATGCAGTGATA
GTAACAATCCGCTAAGAGCGTGGCTCGCATGCTCCGAAAAGCCGCCAGACATATATGC
GATAGGTCTTCAGGAGCTAGACACCCCGACCAAGGCTATGCTCAATAGTACCCAAGTCC
AAGCAATTGAAAAACAGTGGATAGATAAGATGATGGACAGTGTGCATCCTGACGTAGAG
TACGAGATTTTGATGTCTCACCGATTAGTTGCGACGATGCTTACTGTCATCGTTAGAAAA
CAGCTTAGGCAGCACATTATACGTTGTCGACCCAAATCCGTGGCCCGTGGAATATTCAA
CACGCTCGGAAATAAGGGAGGGGTGGCAATATCCCTGCAGTTGAATGAGGGAAACATA
TGTTTTGTTAACTCCCATCTGGCTGCACATATGGGGTACGTCGAAGAGAGGAATCAAGA
TTACAACGCTATTGTCGAAGGCATCAGGTTTGACGACGGTAGAACTATCTCCGACCATG
ACCATATATTTTGGGTAGGAGACTTGAATTATCGAATACAGGAACCTCCGGGACAGCAG
CGTCCTGGGCCGTTGAGTGACGCACAGACATACGAGCTCTTATTACAATACGATCAACT
CCGTCAAGAGATGCGAAGGGGAAAATGCTTCGAAGGTTACACGGAGGGGGAGATTAAG
TTCAGACCGACATACAAATATGATCCTGGAACAGACAATTATGACTCTTCAGAGAAACAG
CGAGCACCTGCATACTGCGATAGAGTGCTATGGAAAGGTACACGTATCGAACAGCTAG
CATACAACAGCATAATGGAAATTAGGCAAAGCGATCATAAGCCAGTTTATGCGGTTTTCC
AGGTGAAGGTAAAGACACGAGATGAGGTGAAGTATAAGCGTGTTCAAGAGGAGGTGCT
AAAGGCGGTTGACAAAAGGGAGAACGATAATCAGCCACAGTAAagcggccgcgaattcgag

**Supplementary Table II**

Synthetic DNA and fluorescent protein coding sequences sequences
